# Lipid-based, protein-based, and steric interactions synergize to facilitate transmembrane signaling stimulated by antigen-clustering of IgE receptors

**DOI:** 10.1101/2020.12.26.424347

**Authors:** Nirmalya Bag, Alice Wagenknecht-Wiesner, Allan Lee, Sophia Shi, David A. Holowka, Barbara A. Baird

## Abstract

Antigen (Ag) crosslinking of immunoglobulin E-receptor (IgE-FcεRI) complexes in mast cells stimulates transmembrane (TM) signaling, requiring phosphorylation of the clustered FcεRI by lipid-anchored Lyn tyrosine kinase. Previous studies showed that this stimulated coupling between Lyn and FcεRI occurs in liquid ordered (Lo)-like nanodomains of the plasma membrane and that Lyn binds directly to cytosolic segments of FcεRI that it initially phosphorylates for amplified activity. Net phosphorylation above a non-functional threshold is achieved in the stimulated state, but not in the resting state, and current evidence supports the hypothesis that this relies on disruption by Ag-crosslinking of a balance between Lyn and tyrosine phosphatase activities. However, the structural interactions that underlie the stimulation process remain poorly defined. This study evaluates the relative contributions and functional importance of different types of interactions leading to supra-threshold phosphorylation of Ag-crosslinked IgE-FcεRI in live rat basophilic leukemia (RBL) mast cells. Our high-precision diffusion measurements by Imaging Fluorescence Correlation Spectroscopy (ImFCS) on multiple structural variants of Lyn and other lipid-anchored probes confirm subtle, stimulated stabilization of the Lo-like nanodomains and concomitant sharpening of segregation from liquid-disordered (Ld)-like regions. With other structural variants we determine that lipid-based interactions are essential for access by Lyn leading to phosphorylation of and protein-based binding to clustered FcεRI. By contrast, TM tyrosine phosphatase, PTPα, is excluded from these regions by steric repulsion of TM segments and preference for Ld-like regions. Overall, we establish a synergy of lipid-based, protein-based, and steric interactions underlying functional TM signaling in mast cells.

**SIGNIFICANCE STATEMENT:** Lipid organization of the plasma membrane is known to be important for facilitating protein interactions in transmembrane signaling. However, the orchestration of these interactions in live cells has been elusive. We employed ImFCS to systemically investigate the interplay of lipids and proteins during signaling in mast cells, initiated as phosphorylation of Ag-crosslinked IgE-FcεRI by lipid-anchored Lyn kinase. We find lipid-based interactions are first required for protein-based phosphorylation of the clustered FcεRI within Lo-like nanodomains. Transmembrane phosphatases must be excluded from these regions, and we find this is mediated by their preference for Ld-like regions and by steric exclusion from the clustered FcεRI proteins. ImFCS provides quantitative characterization of the functional link between features of plasma membrane organization and transmembrane signaling.

## INTRODUCTION

Transmembrane (TM) signaling stimulated by antigen (Ag) occurs through cell surface immunoreceptors that lack a cytosolic kinase module and require tyrosine phosphorylation that is mediated by intermolecular coupling with a separate, plasma membrane localized kinase (1). Effective coupling corresponds to a supra-threshold level of receptor phosphorylation that surmounts dephosphorylation by proximal tyrosine phosphatases. Orchestrated modulation of interactions among the signaling proteins (i.e, receptor, kinase, and phosphatase) (2, 3) and with other proteins (e.g., actin cytoskeleton (4)) are key to Ag-stimulated TM signaling. Although signaling studies have tended to focus on protein-protein interactions, contributions by lipid-based interactions are increasingly appreciated. In particular, phase-like organization of the plasma membrane provides capacity for co-localizing receptor and kinase while segregating phosphatase, according to phase preferences (5). However, the relative importance remains a subject of debate (6–11), largely because it is experimentally difficult to separate the signaling contributions of lipid-based interactions from those of protein-based interactions in live cells.

Our group has worked to develop biophysical approaches that systematically delineate signaling interactions in the context of a prototypical immunoreceptor signaling system: the high affinity receptor for immunoglobulin E (IgE), FcεRI, in RBL mast cells (12, 13). Crosslinking of IgE-FcεRI by soluble, multivalent Ag creates FcεRI nanoclusters (14, 15) that are phosphorylated by Lyn, a src family tyrosine kinase anchored to the inner leaflet of the plasma membrane by saturated acyl chains (Figure 1A). Phosphorylated tyrosines on cytosolic β and γ subunits of FcεRI provide direct binding sites for Lyn-SH2 modules to amplify the phosphorylation activity and create a binding site for Syk kinase and consequent assembly of a protein-based signaling platform that incorporates LAT scaffold and links to activation of phospholipase Cγ and attachment to the actin cytoskeleton (16). In mast cells, stimulated coupling of clustered FcεRI with Lyn initiates the cascade of cellular signaling and responses that underlie allergy and inflammation (12).

**Figure 1.**
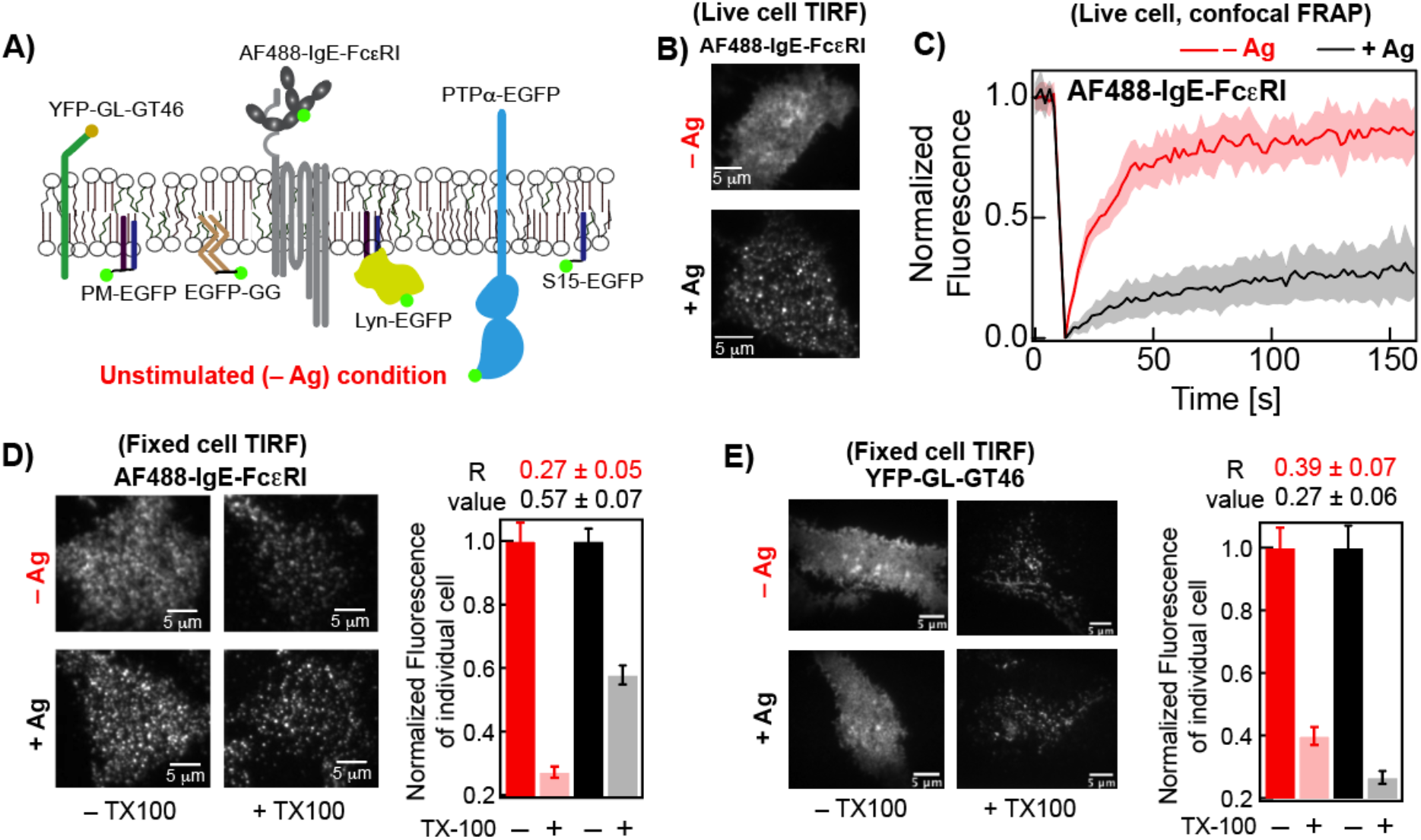
AF488-IgE-FcεRI is clustered, partially immobilized, and exhibits elevated detergent resistance after crosslinking by soluble Ag (DNP-BSA) in RBL cells. **A**) Plasma membrane localization in resting cells (– Ag) of AF488-IgE-FcεRI and other probes evaluated in this study. **B**) Representative TIRF images of AF488-IgE-FcεRI on the ventral plasma membrane in live cells before (−Ag) and after (+ Ag) stimulation by Ag. **C**) Normalized FRAP curves of AF488-IgE-FcεRI obtained from individual cells are overlaid in – Ag (pink) and + Ag (grey) conditions. The solid red and black curves are average of the pink and grey curves, respectively. Figure S1A shows representative fitted FRAP data and box plots of recovery time and mobile fraction of all cells evaluated. **D**) Representative fixed cell TIRF images of AF488-IgE-FcεRI without (–TX100) and with (+ 0.04% TX100) treatment of both – Ag and + Ag conditions. Fluorescence retained after +TX100 treatment is normalized against corresponding –TX100 sample. The *R* values, corresponding to level of detergent resistance, are calculated from the ratio of median fluorescence of multiple cells in +TX100 to –TX100 samples (SI, Eqn S2). The error of *R* values was determined by bootstrapping as described in SI. **E**) Representative fixed cell TIRF images under −/+ Ag and −/+ TX100 conditions and *R* values for YFP-GL-GT46. For each condition, 60-90 cells were imaged from at least two independent sample preparations for both AF488-IgE-FcεRI and YFP-GL-GT46. Box plots of fluorescence values for individual cells under −/+ Ag and −/+ TX100 conditions for both probes in representative experiments are provided in Figure S2.

The role of lipids in stimulated Lyn/IgE-FcεRI coupling has been scrutinized in experimental (15, 17–19) and theoretical (20) studies over two decades, yielding the view that Ag-crosslinking stabilizes liquid ordered (Lo)-like nanodomains around the clustered IgE-FcεRI. Referred to commonly (and roughly) as “rafts,” these proteo-lipid organizational features of plasma membranes have variable properties that depend on cellular circumstance but generally resemble Lo domains that co-exist with liquid disordered (Ld) regions at equilibrium in model membranes of defined composition (21). Experiments in cells show that Lyn kinase preferentially partitions into Lo-like nanodomains in the plasma membrane as mediated by its saturated lipid anchor (palmitoyl/myristoyl, PM) (15, 22, 23). By contrast, a TM tyrosine phosphatase, PTPα, and a Ld-preferring lipid anchor (geranyl/geranyl, GG) preferentially localize to Ld-like regions, away from Lo-like nanodomains (15, 23, 24). In early studies, the functional relevance of this lipid-driven spatial partitioning of the signaling components was shown by appearance of phosphorylated IgE-FcεRI in the Lo-like, detergent resistant membrane (DRM) fraction only after Ag-stimulation (24–26). Further functional evidence came from showing abrogated FcεRI phosphorylation after pharmacological depletion of cholesterol, a key component of Lo-like nanodomain formation (26). However, both DRM isolation and cholesterol depletion have known limitations (27) and they do not directly show whether lipid-based partitioning of Lyn is necessary or sufficient for functional coupling with clustered FcεRI.

Co-clustering of Lyn (and a PM lipid probe, but not a GG lipid probe) with Ag-crosslinked IgE-FcεRI can be observed with fluorescence microscopy but requires pair cross-correlation analysis of super-resolution images and is relatively modest in magnitude. The difficulty of detection points to both lipid-based and protein-based interactions being weak and dynamic. Furthermore, the difference between resting and stimulated state organization of the plasma membrane appears to be subtle, also difficult to detect by conventional fluorescence microscopy and spectroscopy (15, 28, 29). Single particle tracking (SPT) can be successful in picking up small changes. For example, Kusumi and colleagues developed high-speed SPT analysis built around microscope dedicated to delineating transient interactions at single molecule level (30, 31). Both super-resolution and SPT approaches are technically demanding and typically require special fluorescent tags. As a complementary approach, we recently demonstrated that Imaging Fluorescence Correlation Spectroscopy (ImFCS) quantifies subtle differences in the diffusion properties of comparable probes and subtle changes in the diffusion of a particular probe under different cell treatments. ImFCS images conventional fluorophores with a diffraction-limited total internal reflection fluorescence (TIRF) microscope (32, 33). These measurements simultaneously evaluate probe diffusion in hundreds of pixel units, which can be further extended over multiple cells, yielding thousands of data points. Such large data statistics permit precise evaluation of diffusion coefficients thereby enabling detection of small changes in diffusion of a given probe that arises from the change in membrane organization. For example, by comparing Lo-preferring and Ld-preferring probes, we previously detected changes in plasma membrane phase-like organization after inhibition of actin polymerization (33).

Faced the challenge of delineating contributions of protein-based and lipid-based interaction and subtle changes that occur in the plasma membrane after Ag-crosslinking of IgE-FcεRI, we have now improved further the robustness of ImFCS data analysis. We combine with Fluorescence Recovery after Photobleaching (FRAP) and a modified, image-based DRM assay (34), to compare diffusion properties of multiple probes which, as a composite, represent membrane organization under specified conditions. We compare probes for signaling components IgE-FcεRI, Lyn, and PTPα as well as structural variants of these (Figure 1A), and we compare resting and Ag-stimulated steady-states of RBL cells. By evaluating diffusion properties of passive lipid probes with variable Lo-preference we characterize a relatively stable Lo-like environment around crosslinked FcεRI nanoclusters in the stimulated steady-state. We construct Lyn variants to modify Lyn’s lipid-based partitioning, cytosolic protein-based interactions, and kinase activity. Comparative evaluation of these variants reveals levels of contribution toward effective coupling with Ag-crosslinked FcεRI. We show that Lyn’s Lo-preference is essential for stimulated phosphorylation of IgE-FcεRI in a reconstituted system. In contrast, we find that the tyrosine phosphatase, PTPα, is excluded from the region of clustered FcεRI both because of its inherent Ld-preference and by steric hindrance of its TM segments. Overall, this study provides key experimental evidence to explain how cells utilize subtle changes in the membrane organization to initiate TM signaling.

## RESULTS

Ag-crosslinking of IgE-FcεRI in RBL mast cells stimulates new interactions within the plasma membrane to facilitate supra-threshold FcεRI phosphorylation by Lyn tyrosine kinase as the first step of transmembrane (TM) signaling. To delineate comprehensively functional redistributions of lipids and proteins we use multiple complementary approaches to monitor Ag-stimulated changes in diffusion and other properties of selected probes (Figure 1A and following figures). We find that shifts in distribution curves of diffusion coefficients derived from ImFCS provide an exceptionally sensitive representation of membrane changes that occur to initiate signaling.

### ImFCS readily detects subtle changes in lipid heterogeneity caused by IgE-FcεRI clustering

Crosslinking of AlexaFluor488-labeled IgE-FcεRI (AF488-IgE-FcεRI; Figure 1A) by Ag (DNP-BSA) in RBL plasma membranes forms densely distributed nanoclusters that are visible by diffraction-limited TIRF microscopy (Figure 1B). Our previous super-resolution imaging showed that these individual nanoclusters have an average radius of ~80 nm (14). As previously measured by FRAP (35, 36) and confirmed here, 70% of the crosslinked AF488-IgE-FcεRI are immobile, and the 30% mobile fraction diffuse slower (i.e., longer fluorescence recovery time) than monomeric AF488-IgE-FcεRI (85% mobile) present in resting cells (Figure 1C and SI Figure S1A). By comparison, the yellow fluorescent protein (YFP) tagged Ld-preferring probe, YFP-GL-GT46 (comprising the TM segment of the LDL receptor and the cytoplasmic tail of CD46) (37–39) has a mobile fraction of 84% with an insignificant shifts in that value of the fluorescence recovery time after Ag-crosslinking of IgE-FcεRI (Figure S1B).

Changes in resistance to detergent solubilization provided some of the first evidence that crosslinked IgE-FcεRI nanoclusters associate with and stabilize Lo-like nanodomains (5, 15, 22, 24), as similarly documented for B and T cell receptors (34, 40–43). Detergent resistant membranes (DRMs) are Lo-like in lipid composition (44, 45) and retain co-associating proteins after solubilizing cells with 0.04% Triton X 100 (TX100) and floating on sucrose gradients (25). We recently adapted this basic methodology for evaluation of single cells by fluorescence microscopy (46), and we quantify the detergent-resistance of a particular probe in the plasma membrane by a characteristic retention (*R*) value (SI). *R* is taken as the ratio of median fluorescence per cell after treatment with 0.04% TX100 to that of untreated cells. A larger *R* value reflects stronger interaction of the probe with membrane constituents that are not released under these conditions. We find that the *R* value of AF488-IgE-FcεRI increases from 0.27 to 0.57 after antigen crosslinking (Figure 1D and S2A). By comparison, the Ld-preferring TM probe YFP-GL-GT46 shows a slightly smaller *R* value after crosslinking IgE-FcεRI (0.39 to 0.27; Figure 1E and S2B). These distinctive behaviors are consistent with crosslinked AF488-IgE-FcεRI stabilizing Lo-like regions.

We recently established that diffusion properties of mobile membrane probes, and their subtle changes after pharmacological treatments, can be quantified precisely by TIRF-based ImFCS (33). This approach uses a fast camera and autocorrelates fluorescence fluctuations to determine diffusion coefficients (*D*) of a particular probe at several hundreds of diffraction-limited spatial locations (Px unit = 320 nm×320 nm) in single cells (Figure 2A,B). The *D* value determined for an individual Px unit represents all nanoscopic environments that the probe moves through, within that Px unit, during the data acquisition time (280 sec). Nanoscopic environments with more interactions (e.g., higher effective viscosity) yield a slower *D* value, and less interactive environments yield a faster *D* value (Figure 2B,C). By reflecting the interactions that structurally distinct probes experience, distinctive diffusion properties provide information about membrane organization. Pooling data from multiple cells, yields ~10,000 *D* values for a given probe, and the resulting high level of precision enables subtle differences in diffusion properties among probes to be discerned ((33); Table S1). As one measure of precision, the arithmetic average of pooled *D* values, *D*_av_, has a standard error of the mean (SEM) of less than 1 % for every probe evaluated by ImFCS in this study.

**Figure 2.**
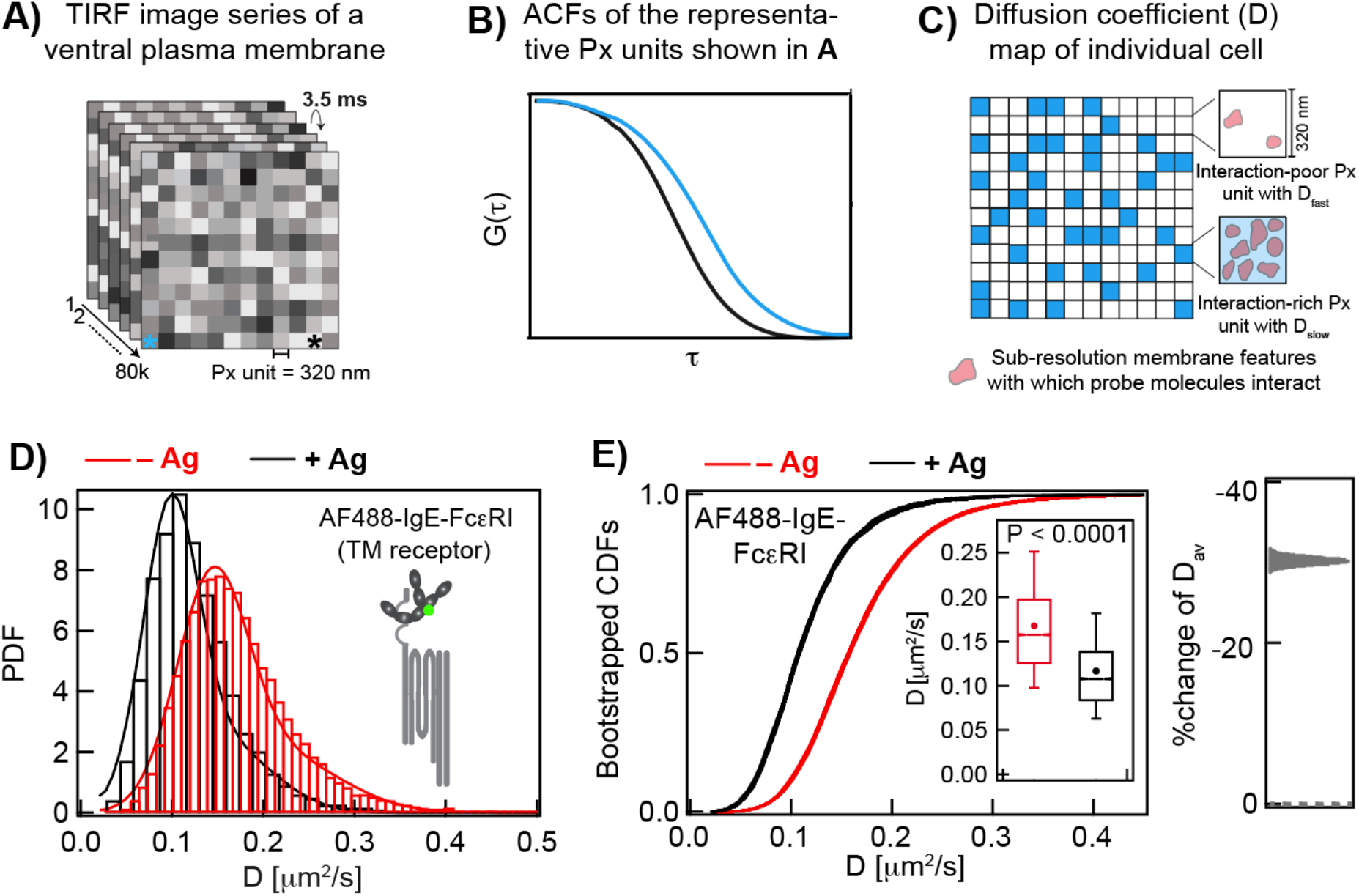
Large data sets of ImFCS precisely characterize spatially heterogeneous diffusion of plasma membrane probes in both unstimulated (−Ag) and stimulated (+ Ag) cells. **A-B**) In a typical ImFCS recording, 80,000 TIRFM images of fluorescently labeled ventral plasma membrane are collected at 3.5 ms/frame. The autocorrelation function (ACF) from a given Px unit (320 nm × 320 nm) decays faster if probes diffuse faster within that Px unit. The ACFs, corresponding to the Px units designated with asterisks of same color, illustrate probes diffusing slower (cyan) and faster (black). **C**) Schematic diffusion coefficient (*D*) map, obtained after ACF analyses of all Px units, contains some Px units with relatively slower (*D*_slow_, cyan) or faster (*D*_fast_, white) diffusion coefficient, interpreted as interaction-rich and interaction-poor units, respectively. **D)** Histograms of experimental *D* values (>10,000; SI Table S1) and probability distribution function (PDF) for AF488-IgE-FcεRI at −Ag (red) and +Ag (black) steady-states. PDFs are fitted using parameters derived from bin-independent cumulative distribution functions (CDFs; SI Eqn A2). **E**) CDFs of the same *D* values as in part (D). Pooled *D* values are resampled 30 times by bootstrapping with 50% of all data each time and individual bootstrapped CDFs are fitted for *D*_slow_, *D*_fast_, and F_slow_ (SI Appendix; Table S1 and Figure S3). Individual raw bootstrapped CDFs of *D* values of AF488-IgE-FcεRI at each condition are overlaid and shown (red: −Ag and black: +Ag). Inset: Box plots of all *D* values. Box height corresponds to 25^th^ to 75^th^ percentile; error bars represent 9^th^ to 91^st^ percentile of entire data set; mean and median values are represented as solid circle and bar, respectively; notches signify 95% confidence interval of the median. Stimulated %change of *D*_av_: Distribution is calculated from the bootstrapped mean values at each condition.

The probability distribution functions (PDFs) of *D* values determined by ImFCS for mobile AF488-IgE-FcεRI from resting cells (red) and from cells stimulated with antigen (black) show a clear shift to slower *D* values (Figure 2D), consistent with FRAP measurements (Figures 1C, S1A). To quantify subpopulations we convert the pooled *D* values into cumulative distribution functions (CDFs; Figure 2E) – a mathematically equivalent but bin-independent alternative to PDFs – which can be unequivocally resolved into one or two Gaussian components (SI Eqns S3 and S4; (33)). For two-component *D* CDFs, the fast and slow components, *D*_fast_ and *D*_slow_ represent the average diffusion coefficient of a particular probe in, respectively, interaction-poor and interaction-rich populations of Px units in the plasma membrane (Figure 2C). *F*_slow_ is the fraction of the interaction-rich population and (1 – *F*_slow_) is the fraction of the interaction-poor population. To test whether CDF curves are overly influenced by outliers, and to provide a curve thickness related to level of uncertainty, we routinely resample the *D* values by bootstrapping 30 times (with 50% of all data each time), and the corresponding bootstrapped CDFs are fitted individually (See SI for more details of the analysis). For all probes tested in this study we found narrowly distributed fitted values of *D*_fast_, *D*_slow_, and *F*_slow_, confirming the reliability and robustness of our analysis (Table S1; Figures 2–6 and S3).

After Ag-crosslinking AF488-IgE-FcεRI to form nanoclusters (14, 47), ImFCS measurements show the mobile fraction (30% as measured by FRAP; Figure 1C) has ~35% smaller *D*_av_ than monomeric AF488-IgE-FcεRI in resting cells (Figure 2D,E; Table S1). Mobile IgE-FcεRI in the stimulated steady state are likely to be small oligomers, which have been shown to be signaling-competent (48). These oligomers diffuse through interaction-poor and -rich populations of Px units with ~30% reduction in both *D*_fast_ and *D*_slow_, and unchanged *F*_slow_ after stimulation, compared to the resting steady state (Table S1). It is also possible that immobilized IgE-FcεRI clusters reduce effective membrane area accessible to mobile IgE-FcεRI oligomers and other TM proteins, thereby restricting diffusion of the mobile TM species.

The CDF curves of *D* values measured with ImFCS and precise fitted values of diffusion parameters (*D*_av_, *D*_fast_, *D*_slow_, *F*_slow_) were determined for all the probes evaluated in this study (Table S1 and Figure S3). These parameters quantify how structurally distinct probes sense changes in local environments caused by Ag-mediated clustering of IgE-FcεRI to stimulate TM signaling. Visually striking are the distinctive shifts in the CDF curves that accompanies this stimulation (Figures 2 – 6); these shifts represent changes in interactions for a given probe after stimulation. Comparing such curve shifts for a panel of probes with defined structural features allows us to evaluate contributions of individual structural features to changes in diffusion properties. In this manner we can infer how stimulation changes lipid-based or protein-based interactions in the plasma membrane and correspondingly changes in membrane organization.

To focus first on changes occurring in lipid phase-like properties and the Lo-like environment stabilized by clustered IgE-FcεRI, we evaluated lipid probes that are phase-selective, but otherwise passive in the signaling process. We employed three established inner leaflet lipid probes (Figure 1A; (23, 49)): PM-EGFP (Lo-preferring; (23)), EGFP-GG (Ld-preferring; (23)) and S15-EGFP (Ld-preferring; (50)). Our previous super-resolution imaging demonstrated that the PM construct co-clusters with crosslinked IgE-FcεRI but that the GG construct does not (15), and consistent results were obtained with clustered B cell receptors (41). However, diffraction-limited TIRF imaging reveals no visible changes in the distribution of any of these probes (Figure S6). We examined their membrane interactions using DRM imaging, FRAP, and ImFCS.

For lipid probes, a larger *R* value in the DRM assay indicates stronger partitioning into Lo-like nanodomains. In resting cells, we found *R* values to be consistent with their phase preferences in membranes as reported previously (49–51): PM-EGFP > EGFP-GG ≈ S15-EGFP (Figure 3A and S4, red/pink). Although these *R* values are useful for monitoring Lo-vs Ld-phase preference, we found the method to be insufficiently sensitive for detecting significant differences before and after crosslinking IgE-FcεRI by Ag (Figure 3A and S4, compare red/pink to black/gray). Similarly, FRAP measurements (Figure 3C, E, G and S5A-C) do not resolve significant differences among these probes before and after stimulation. In contrast, Ag-crosslinking of IgE-FcεRI causes distinctive shifts in the ImFCS CDF curves of these probes (Figure 3B,D,F): Lo-preferring probe PM-EGFP shifts to slower *D* values, and Ld-preferring probes EGFP-GG and S15-EGFP shift to faster *D* values. In terms of *D*_av_ values, the shifts are PM-EGFP (0.62 to 0.57 μm^2^/sec; 8% slower), EGFP-GG (0.64 to 0.69 μm^2^/sec; 8% faster), S15 (0.90 to 0.92 μm^2^/sec; 2% faster). This opposing behavior of Lo- and Ld-preferring lipid probes is consistent with the Lo-like phase in the inner leaflet of the plasma membrane becoming more ordered and stabilized against surrounding Ld-like regions after Ag addition. Fitting these curves with one (S15-EGFP) or two Gaussian populations yields distinctive values for *D_fast_, D_slow_*, and *F_slow_* for each of these probes (Table S1), further revealing how they differentially sense the same membrane milieu and providing additional insight into stimulated changes in membrane organization. These ImFCS measurements on inner leaflet lipid probes are consistent with and extend quantitative results from previous super-resolution imaging (15) and high-speed SPT (30). Collectively, they characterize and quantify stabilization of an Lo-like environment encompassing Ag-crosslinked IgE-FcεRI nanoclusters.

**Figure 3.**
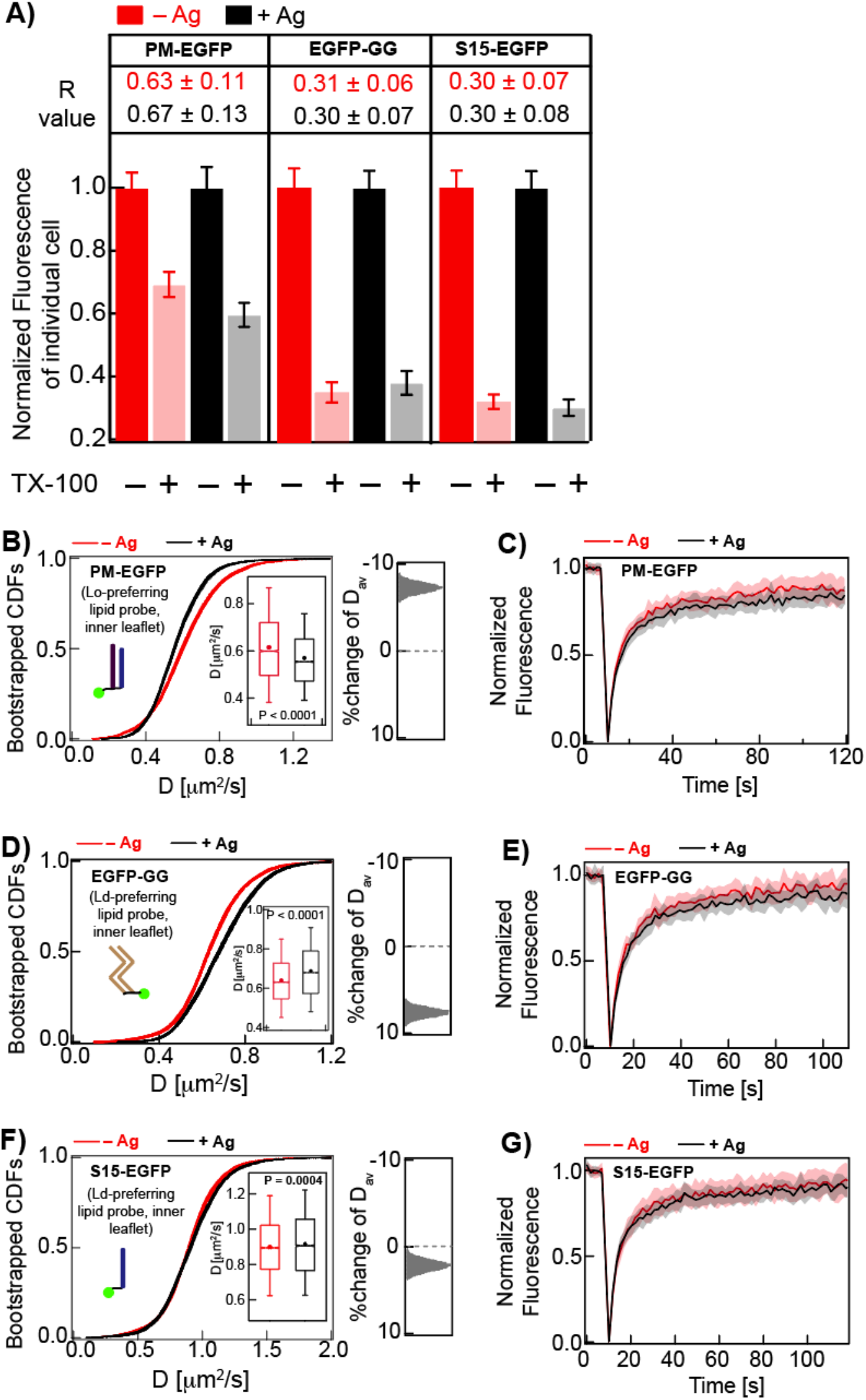
ImFCS, but not DRM and FRAP, detects subtle stabilization of Lo-like nanodomain in Ag-stimulated RBL cells. **A**) Degree of detergent resistance for PM-EGFP, EGFP-GG, and S15-EGFP and *R* values for –Ag (red) and +Ag (black) conditions. Box plots of fluorescence values of individual cells for −/+ TX100 and −/+ Ag conditions for these probes are provided in Figure S4A-C. **B, D, F)** 30 bootstrapped CDFs of *D* values from ImFCS measurements are overlaid for specified probes and conditions (−/+ Ag). Box plots of all *D* values and distribution of stimulated %change of *D*_av_ as described for Figure 2E. Table S1 shows number of ACF and cells measured for ImFCS analyses. **C, E, G)** Normalized FRAP curves for specified probe obtained from many cells are overlaid for – Ag (pink) and + Ag (grey) conditions. Figure S5A-C show representative fitted FRAP data and box plots of recovery time and mobile fraction for all cells.

### Lo-preference of lipid anchor is required for Lyn’s initial coupling with crosslinked IgE-FcεRI

Nanoscopic co-localization of Lyn and crosslinked IgE-FcεRI nanoclusters is measurable but weak, as demonstrated super-resolution imaging (15). This weak stimulated modulation in Lyn interactions is not detectable in TIRF images (Figure S7A), DRM *R* values (Figure S7B), or by FRAP curves (Figures 4C and S7C). However, ImFCS is more sensitive and detects the change by resolving a shift of the CDF curves to lower *D* values (Figure 4B), corresponding to 10% reduction of *D*_av_ of Lyn-EGFP after antigen addition (from 0.49 to 0.44 μm^2^/s; Table S1).

**Figure 4.**
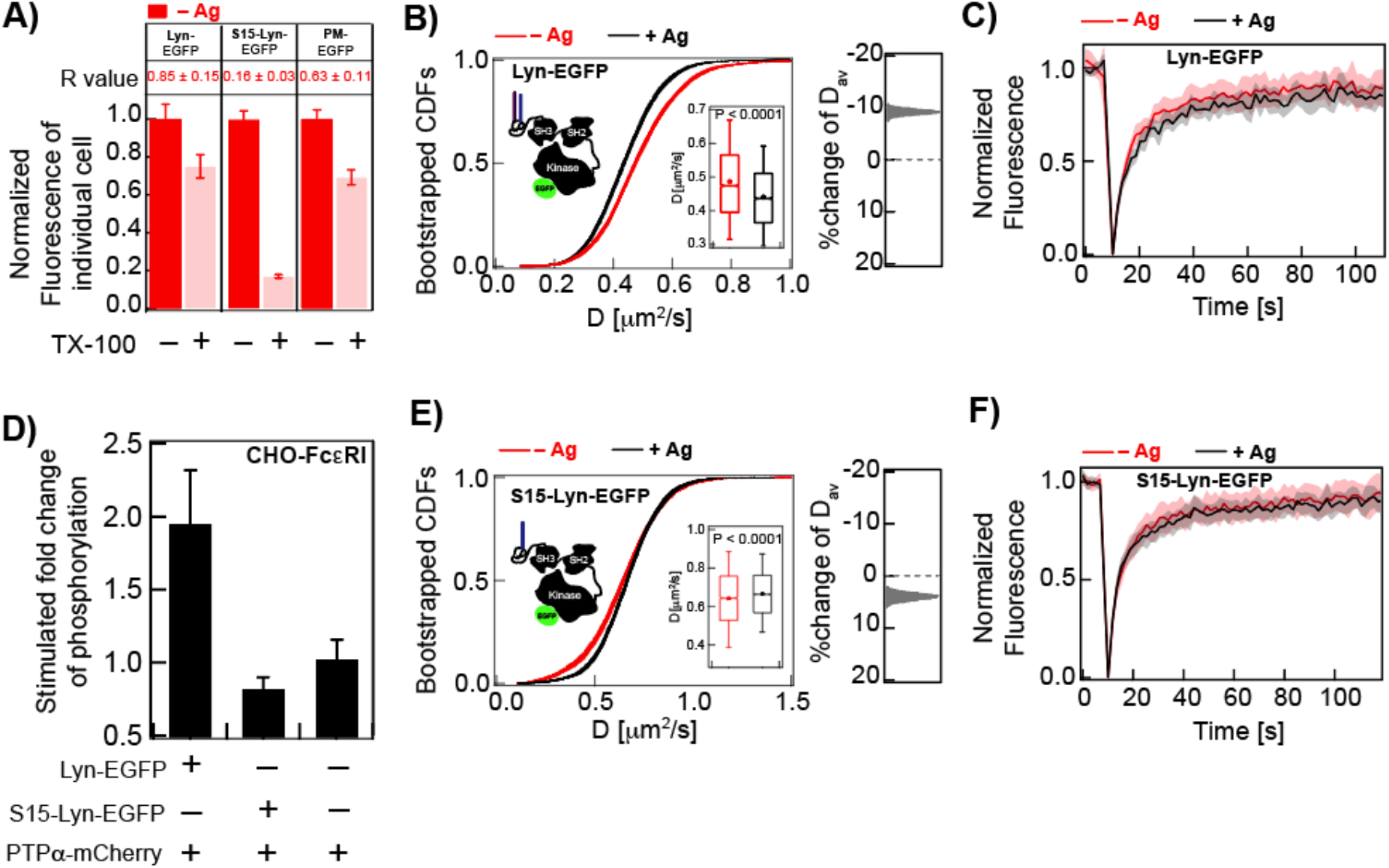
Lipid-driven Lo-preference of Lyn is necessary for functional coupling with Ag-crosslinked IgE-FcεRI. **A)** Detergent resistance of Lyn-EGFP, S15-Lyn-EGFP, and PM-EGFP represented by relative loss of fluorescence after TX100 treatment and corresponding *R* values in unstimulated (−Ag) RBL cells. Box plots of fluorescence values of individual cells under −/+ TX100 and −/+ Ag conditions are provided in Figures S4A (PM-EGFP) and S7B (Lyn-EGFP and S15-Lyn-EGFP). **B, E)** 30 bootstrapped CDFs of *D* values from ImFCS measurements are overlaid for specified probes and conditions (-/+ Ag). Box plots of all *D* values and stimulated %change of *D*_av_ are shown as described for Figure 2E. Table S1 shows number of ACF and cells measured for ImFCS analyses. **C, F**) Normalized FRAP curves for specified probes and conditions. Figure S7C-D show representative fitted FRAP data and box plots of recovery time and mobile fraction for all cells. **D)** Stimulated fold change of phosphorylation determined from anti-pTyr (4G10) immunostaining of CHO cells stably transfected with FcεRI and transiently transfected with specified Lyn variant and PTPα.

Lyn-EGFP is highly detergent resistant (Figure 4A; (28)), as expected for the Lo-preference of its palmitoyl/myristoyl membrane anchor and consistent with the stimulated shift in this probe’s diffusion properties being the consequence of more stable Lo-like nanodomains after antigen-crosslinking of IgE-FcεRI. However, the shifts in the CDF curves are different for Lyn-EGFP (Figure 4B) compared to PM-EGFP (Figure 3B), as quantified by distinctive values and trends of *D*_av_, *D*_fast_, *D*_slow_, and *F*_slow_ for the two probes (Figure S3; Table S1). These differences indicate that some of Lyn-EGFP stimulated interactions are directly with the crosslinked FcεRI, in addition to those with the surrounding stabilized Lo-like regions into which the PM lipid anchor partitions favorably.

To deconvolve lipid-based and protein-based interactions, we compared diffusion modes of Lyn-EGFP in both resting and stimulated states to a Lyn chimera. S15-Lyn-EGFP was created by replacing the first 15 amino acids of wt Lyn which possesses Lo-targeting palmitoylation and myristoylation sites with the first 15 amino acid of Ld-preferring lipid probe S15-EGFP (50). Unlike Lyn-EGFP and PM-EGFP, S15-Lyn-EGFP is weakly detergent-resistant (Figure 4A) and has a faster *D*_av_ in resting cells (– Ag conditions in Figures 3B and 4B,E; Table S1) and serves as an Ld-preferring Lyn probe that possesses all functional protein modules. If protein-protein interactions primarily drive stimulated reduction of Lyn-EGFP diffusion, we expect to see similar net change of diffusion of S15-Lyn-EGFP after cross-linking IgE-FcεRI. However, the *D*_av_ of S15-Lyn-EGFP (Figure 4E), in sharp contrast to *D*_av_ values for Lyn-EGFP (Figure 4B) and PM-EGFP (Figure 3B), does not decrease after antigen addition. In fact, it increases slightly from 0.64 μm^2^/s to 0.67 μm^2^/s similar to the behavior of Ld-preferring lipid probes, EGFP-GG (Figure 3D) and S15-EGFP (Figure 3F). For direct numerical comparisons of these ImFCS values, see also Table S1 and Figure S3. Notably, data statistics of FRAP are not sufficient to detect these differences (Figures 3C,E,G; 4C,F; S5A-C; and S7C-D). The slower diffusion of Lyn-EGFP and faster diffusion of S15-Lyn-EGFP after crosslinking IgE-FcεRI are revealed by ImFCS, and these reflect a distinctive but subtle change in the phase-like organization that is sensed by Lo- and Ld-preferring probes. These ImFCS results for Lyn-EGFP, PM-EGFP and S15-Lyn-EGFP are consistent with the view that Lo-preference is important for Lyn’s interaction with clustered IgE-FcεRI, and also point to a role for protein-based interactions for optimal coupling leading to receptor phosphorylation and downstream signaling.

### Lyn-EGFP but not S15-Lyn-EGFP facilitates Ag-dependent tyrosine phosphorylation

To test directly a functional outcome inferred from our ImFCS results, we compared the capacities of Lyn-EGFP and S15-Lyn-EGFP to facilitate stimulated phosphorylation in cells. We used a reconstitution approach for this purpose: FcεRI stably expressed in Chinese Hamster Ovary (CHO) cells that are transiently co-transfected with Lyn and an Ld-preferring TM tyrosine phosphatase (PTPα). With this experimental system, we found previously that PTPα suppresses Lyn kinase activity and minimizes its spontaneous phosphorylation of FcεRI, while facilitating Ag-stimulated FcεRI phosphorylation (24). Importantly, an Lo-preferring chimeric version of PTPα failed to reconstitute Ag-dependent FcεRI phosphorylation. In the current study we used immunostaining to monitor tyrosine phosphorylation in CHO cells stably expressing FcεRI and transiently transfected with PTPα and either Lyn-EGFP or S15-Lyn-EGFP, before and after addition of Ag. As shown in Figure 4D, co-transfection with Lyn-EGFP causes a nearly two-fold enhancement of tyrosine phosphorylation, compared to no Lyn construct, whereas co-transfection with S15-Lyn-EGFP results in a somewhat reduced level of tyrosine phosphorylation. Results from this functional assay are consistent with ImFCS diffusion measurements and show that Lo-preference is prominently involved in Lyn’s initial sensing of clustered IgE-FcεRI, leaving open the possibility that protein-based interactions are also involved.

### Intact protein modules of Lyn are necessary for its effective coupling with crosslinked IgE-FcεRI

The cytosolic segments of Lyn comprise an SH2 module that docks on phosphotyrosine (pY) sites, an SH3 module which interacts with polyproline (PxxP) motifs, and a kinase module (52). To test the importance of direct interactions with FcεRI in stimulated cells, we evaluated the diffusion properties of two point mutants (53): Lyn-mSH2-EGFP (Arg to Ala at position 135) and Lyn-mSH3-EGFP (Trp to Ala at position 78), with intact PM lipid anchor but disabled binding capabilities through SH2 and SH3 modules respectively. In resting cells, the detergent-resistance of both mutants (*R*~0.6) is less than Lyn-EGFP (Figure 5A), similar to PM-EGFP (Figure 3A) and greater than Ld-preferring S15-Lyn-EGFP (*R*~0.2; Figure 4A). Changes in diffusion properties of Lyn-mSH2-EGFP and Lyn-mSH3-EGFP after addition of antigen to crosslink IgE-FcεRI are not detected by FRAP (Figure 5C,E and S8B,C) but revealed by ImFCS (Figure 5B,D). Importantly, ImFCS further reveals clear differences for these two Lyn mutants (Figure 5B,D) compared to Lyn-EGFP (Figure 4B) in terms of shifts in CDF curves caused by Ag. This is difference is quantified by the values of *D*_av_, *D*_fast_, *D*_slow_, and *F*_slow_ derived from these curves (Table S1, Figure S3). For example, after antigen addition the *D*_av_ values of Lyn-EGFP decreases (Figure 4B), for Lyn-mSH2-EGFP increases slightly (Figure 5B), and for Lyn-mSH3-EGFP decreases slightly (Figure 5D). This comparison indicates that Lyn-mSH2-EGFP responds differently from PM-EGFP to stimulated stabilization of Lo-like nanodomains, even though it is similarly detergent-resistant. One possible explanation for differences is cytosolic steric hindrance due to scaffold proteins and cytoskeleton components that are recruited to antigen-crosslinked IgE-FcεRI (13, 54, 55). Lo-preferring lipid, PM-EGFP, with no Lyn protein modules in the cytoplasm, is likely to be less sensitive to these steric factors. Unlike Lyn-EGFP, Lyn-mSH2-EGFP and Lyn-mSH3-EGFP cannot efficiently overcome steric hindrance by docking at the pY219 site on clustered FcεRI. It appears that antigen-induced changes in Lyn diffusion compared to these two Lyn mutants depend on interactions with FcεRI rather than entirely on their Lo-preference.

**Figure 5.**
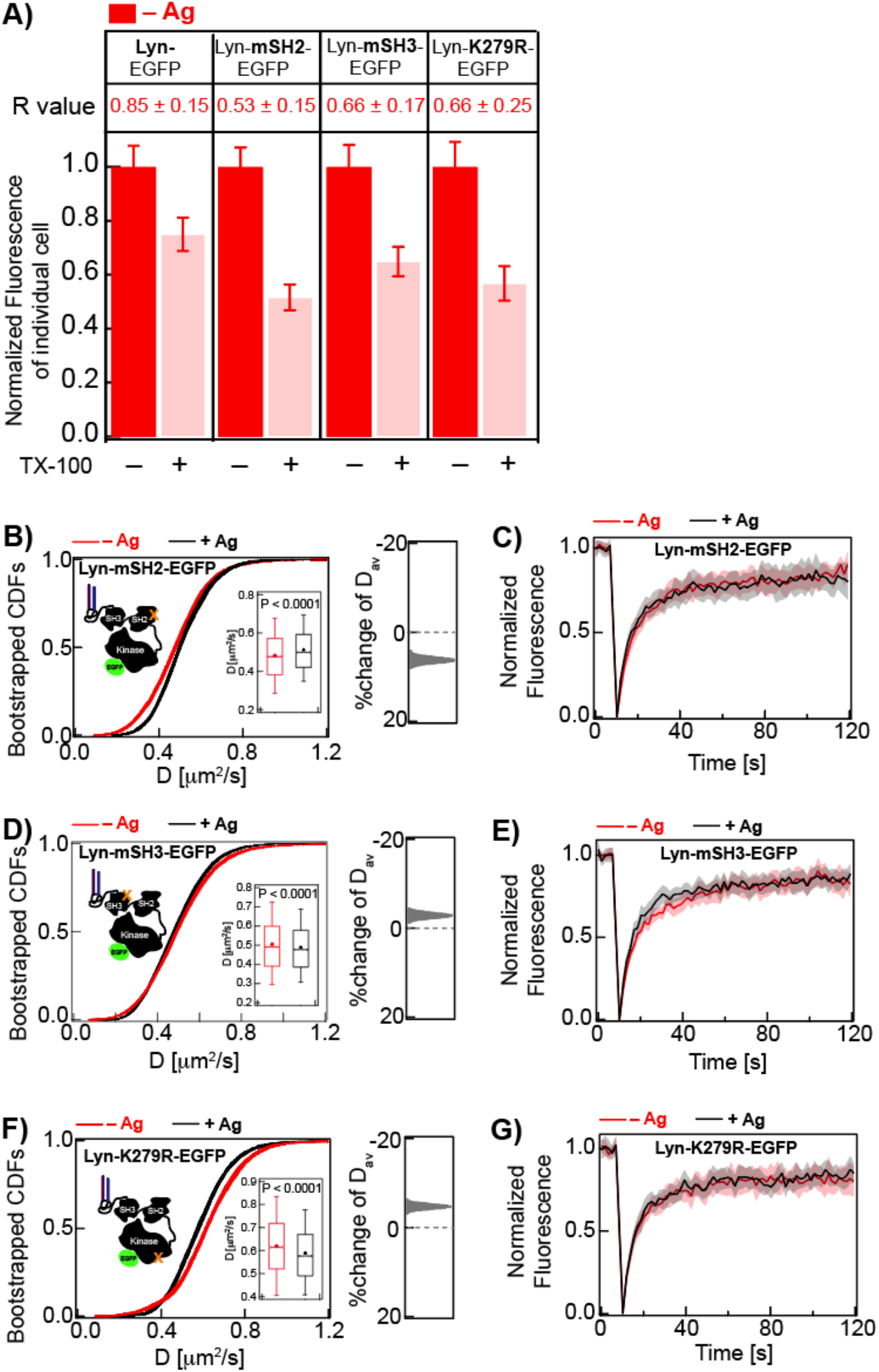
Cytosolic protein modules of Lyn-EGFP contribute to detergent-resistance and reduction of diffusion caused by Ag-crosslinking of IgE-FcεRI. **A)** Detergent resistance of Lyn-EGFP compared to point mutants Lyn mSH2-EGFP, Lyn-mSH3-EGFP, and Lyn-K279R-EGFP as represented by relative loss of fluorescence after TX100 treatment and corresponding *R* values in unstimulated (– Ag) RBL cells. Box plots of fluorescence values of individual cells for −/+ TX100 and −/+ Ag conditions for these probes are provided in Figure S7B (Lyn-EGFP) and S8A (Lyn-EGFP variants). **B, D, F)** 30 bootstrapped CDFs of *D* values from ImFCS measurements are overlaid for specified probes and conditions (−/+ Ag). Box plots of all *D* values and stimulated %change of *D*_av_ are shown as described for Figure 2E. Table S1 shows number of ACF and cells measured for ImFCS analyses. **C, E, G)** Normalized FRAP curves for specified probes and conditions. Figure S8B-D show representative fitted FRAP data and box plots of recovery time and mobile fraction for all cells.

We also measured the stimulated diffusion change of kinase-inactive Lyn mutant, Lyn-K279R-EGFP (56), to test whether phosphorylation of FcεRI by Lyn stabilizes the coupling of these two proteins. In resting cells, similar to Lyn-mSH2-EGFP and Lyn-mSH3-EGFP, Lyn-K279R-EGFP shows weaker detergent-resistance (*R*~0.6) than wt Lyn-EGFP (Figure 5A and S8D). This consistent difference suggests that kinase activity and resulting protein-based interactions contribute to Lyn’s greater tendency to localize in an Lo-like environment. Lyn-K279R-EGFP, unlike Lyn-mSH2-EGFP and Lyn-mSH3-EGFP, can undergo proper SH2/pY219 intermolecular docking after FcεRI phosphorylation by the endogenous Lyn present in these cells (56). However, the shift in the CDF curve to lower diffusion coefficients is clearly less for Lyn-K279R-EGFP (5% decrease of *D*_av_: 0.62 to 0.59 μm^2^/sec; Figure 5F) compared to Lyn-EGFP (10% decrease of *D*_av_: 0.49 to 0.44 μm^2^/sec; Figure 4B), indicating competitive advantage of endogenous Lyn over Lyn-K279R in associating with phosphorylated FcεRI. Again, FRAP failed to detect this modest difference (Figure 5G and S8D).

Collectively, the shifts in CDF curves (Figures 4B and 5B,D,F) and underlying ImFCS parameters (Figure S3; Table S1) of Lyn-EGFP compared to the variants we tested support the following view: the Lo-preference of Lyn, as mediated by saturated lipid anchors, is required for its initial coupling with antigen-crosslinked FcεRI; then Lyn’s kinase activity and FcεRI-docking capacities serve to stabilize the interaction. Lyn variants with mutated modules that lose these capacities may be subject to steric hindrance, resulting in reduced coupling.

### Tyrosine phosphatase PTPα spatially segregates from FcεRI nanoclusters by steric and lipid-based exclusion

Maintaining the phosphorylated state of Ag-crosslinked FcεRI above the stimulation threshold, requires that access by phosphatases be minimized. As previously hypothesized (5, 22, 24, 57), tyrosine phosphorylation of FcεRI by Lyn kinase prior to antigen engagement is counter-balanced by TM phosphatase-mediated dephosphorylation; transient nanodomains present in the resting steady-state do not sufficiently co-confine Lo-preferring Lyn and FcεRI nor prevent access by phosphatase to disrupt the balance. Lo-like nanodomains that are stabilized around the antigen-clustered FcεRI preferentially include kinase, exclude phosphatase, and tip the balance to exceed the phosphorylation threshold. We examined participation of the TM phosphatase within this mechanism by measuring stimulated changes in diffusional and other properties of EGFP-tagged protein tyrosine phosphatase α (PTPα-EGFP) variants.

Consistent with our previous observations that PTPα prefers an Ld-like environment (24), we found that PTPα-EGFP is almost completely detergent-soluble (*R* = 0.1; Figure 6A and S9A). ImFCS measurements of this probe in resting cells yielded *D*_av_ = 0.23 μm^2^/s, which is slower than lipid and lipid-anchored probes, as expected for a TM probe (33, 38). If the diffusion of PTPα-EGFP is influenced only by its Ld-preference, we expect Ag-stimulation to increase its *D*_av_, similar to EGFP-GG or S15-EGFP (Figure 3D,F). However, the *D*_av_ of PTPα-EGFP is reduced by 13% after receptor cross-linking (Figure 6B, Table S1). This decrease and corresponding shift in CDF curves to lower diffusion coefficients (Figure 6B) is similar to that for Lyn-EGFP (Figure 4B), which undergoes direct binding interactions with cytosolic segments of FcεRI. However, PTPα-EGFP lacks cytosolic docking modules (58) and is preferentially excluded from the Lo-like regions stabilized around clustered FcεRI (24). Therefore, some other stimulated change in the environment of PTPα-EGFP apparently causes its reduced diffusion.

**Figure 6.**
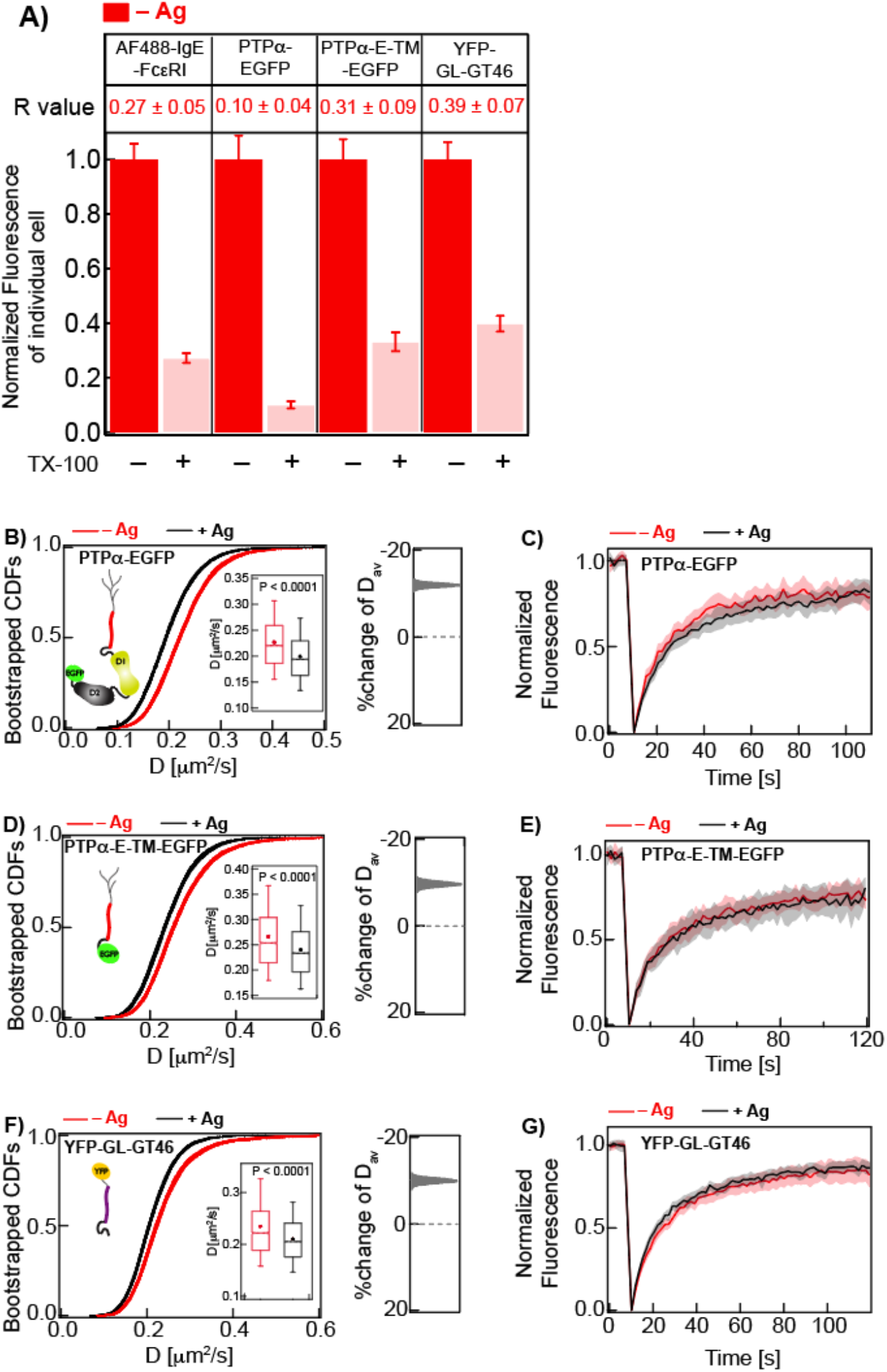
TM probes are strongly detergent-soluble but show relatively slower diffusion in stimulated cells, likely due to steric obstruction by TMDs of Ag-clustered FcεRI. **A)** Detergent resistance of AF488-IgE-FcεRI, PTPα-EGFP, PTPα-E-TM-EGFP, and YFP-Gl-GT46 as represented by relative loss of fluorescence after TX100 treatment and corresponding *R* values in unstimulated (−Ag) RBL cells. Box plots of fluorescence values of individual cells for −/+ TX100 and −/+ Ag conditions for these probes are provided in Figure S2 (AF488-IgE-FcεRI and YFP-GL-GT46) and S9A (PTPα-EGFP and PTPα-E-TM-EGFP). **B, D, F)** 30 bootstrapped CDFs of *D* values from ImFCS measurements are overlaid for specified probes and conditions (-/+ Ag). Box plots of all *D* values and stimulated %change of *D*_av_ for these samples are shown as described for Figure 2E. Table S1 shows number of ACF and cells measured for ImFCS analyses. **C, E, G**) Normalized FRAP curves for specified probes and conditions. Figures S1 and S9B-C show representative fitted FRAP data and box plots of recovery time and mobile fraction for all cells.

We considered the possibility that the IgE-FcεRI nanoclusters immobilized by Ag-crosslinking (70% of total IgE-FcεRI; Figure 1C) impose impermeable diffusion obstacles for TM probes in stimulated cells (59). In other words, these probes’ TM domains (TMDs) are sterically excluded from regions defined by TMDs of the immobilized FcεRI. We expect these regions are much more accessible to lipid-anchored probes, which can diffuse into nanoclusters and explore the lipid membrane among individual TMDs. To test this hypothesis, we created a construct, PTPα-E-TM-EGFP comprising only the extracellular portion and TMD of PTPα and C-terminally fused to EGFP (as illustrated in Figure 6D). Because this probe does not contain the catalytic or other cytosolic modules, any Ag-stimulated change of its diffusion is not due to functional interactions. PTPα-E-TM-EGFP shows strong detergent-solubility and FRAP similar to PTPα-EGFP (Figure 6A,C,E and S9A-C). The *D*_av_ (0.27 μm^2^/s) of PTPα-E-TM-EGFP in resting cells is slightly greater than the PTPα-EGFP (Table S1) possibly due to eliminated cytosolic interactions. After Ag-crosslinking of IgE-FcεRI, the *D*_av_ of PTPα-E-TM-EGFP decreases by ~11%, and the shift in ImFCS CDF curves is strikingly similar to PTPα-EGFP (Figure 6B,D), as quantified by ImFCS parameters (*D*_av_, *D*_fast_, *D*_slow_ and *F*_slow_; Figure S3). This comparison indicates that Ag-stimulated reduction of PTPα diffusion is not due cytosolic protein-based interactions. Instead, the stimulated diffusion reduction of PTPα-EGFP is likely due to its TMD which are blocked by impermeable FcεRI nanoclusters (60). We tested this hypothesis on YFP-GL-GT46, the Ld-preferring TM probe (37–39), with both extracellular and cytosolic domains shorter than those of PTPα-EGFP. As expected, YFP-GL-GT46 is highly detergent-soluble in resting cells (Figure 1E and 6A). After Ag-crosslinking of IgE-FcεRI, this probe exhibits a clear shift in ImFCS CDF curves to lower diffusion coefficients that is very similar to those of PTPα-EGFP and PTPα-E-TM-EGFP (Figure 6B,D,F), as quantified by their respective ImFCS parameters (Figure S3). Together, these measurements support the view that Ag-crosslinked, immobile IgE-FcεRI nanoclusters obstruct diffusion of mobile TM probes (mobile fractions of AF488-IgE-FcεRI, Figure 1C and S1A; PTPα-EGFP, Figure 6C and S9B; PTPα-E-TM-EGFP, Figure 6E and S9C; YFP-GL-GT46, Figure 6G and S1B), independently of respective ectodomain size, cytosolic segments, and detergent-resistance.

## DISCUSSION

Nanoclustering of TM receptors by extracellular ligands followed by local reorganization in the plasma membrane to facilitate kinase coupling and receptor phosphorylation above a stimulation threshold has become a general paradigm of signaling through cell surface immunoreceptors. A large body of evidence supports the view that Ag-mediated nanoclustering of sensitized mast cell receptors, IgE-FcεRI, coalesces Lo-like nanodomains (5, 6, 61). However, the functional significance of lipid phase-like behavior in TM signaling mediated by this and other immunoreceptors continues to be debated (21, 62, 63), and some have argued that stimulated protein-protein interactions are sufficient (3). Although protein-based interactions may increasingly dominate as signaling proceeds, the initial upshift in receptor phosphorylation appears to depend on disrupting the balance between kinase and phosphatase access, and this is mediated primarily by membrane lipids. Simply, stabilization of Lo-like nanodomains around Ag-crosslinked immunoreceptors serves to co-localize Lo-preferring lipid-anchored kinase (e.g., Lyn) while excluding Ld-preferring TM phosphatases (e.g., PTPα). In this study we used ImFCS and other imaging approaches to quantify systematically the subtle shift in plasma membrane organization that accompanies Ag-mediated stimulation. We confirm the primacy of lipid-based interactions and unveil how these synergize with protein-based, and steric interactions among IgE-FcεRI, Lyn kinase, and PTPα phosphatase. We evaluate particular contributions of these different types of interactions for successful assembly of TM signaling components.

We previously established ImFCS as a statistically robust approach for quantifying subtle differences in the diffusion properties of structurally distinct probes that collectively sense the organization of the plasma membrane under a specified condition (33). The sampling provided by our ImFCS measurements (~10,000 *D* data points for each probe) yields extremely precise values for *D_av_*. The CDF of *D* values for each probe are also highly precise, as demonstrated by bootstrapping, and fit parameters *D*_fast_, *D*_slow_, and *F*_slow_, further quantify the membrane environment as it is experienced by a particular probe (33). As we demonstrate herein, ImFCS measurements on 13 independent probes are highly sensitive to subtle changes in plasma membrane organization that result from Ag-stimulation. These changes are quantified by detailed changes in fit parameters (Table S1), simply represented by *D_av_* values and visually obvious in CDF curve shifts (Figures 3 – 6). By systematically changing key structural features, we can reconstruct participation of different types of interactions experienced by the initial signaling components.

### Ag crosslinking creates nanoclusters of IgE-FcεRI and stabilizes surrounding Lo-like membrane domains

The phase-like behavior of the plasma membrane in resting cells is represented by the tendency of probes with saturated lipid anchors (PM-EGFP, Lyn-EGFP) to be more detergent-resistant (Lo-preferring) than those with unsaturated lipid anchors (EGFP-GG, S15-EGFP, S15-EGFP (Ld-preferring) (Figures 3A and 4A). However, the Lo-like domains in this resting state appear to be small and transient (21) with relative weak distinction from Ld-like regions, as suggested by small differences in *D_av_* values for PM-EGFP and EGFP-GG ((33); Figure 3). This relatively weak phase-like heterogeneity in the cell’s resting state would allow access to FcεRI by both Lyn kinase and TM phosphatase and limit net phosphorylation (64). That Ag crosslinking of IgE-FcεRI leads to proximal coalescence and stabilization of Lo-like domains was clearly indicated by functional studies (24) and by super-resolution imaging of co-clustering with IgE-FcεRI by PM probes but not GG probes (15). However, this stabilization is not detected by DRM imaging and FRAP measurements (Figure 3). In contrast, ImFCS measurements show subtle but distinctive shifts in *D* CDF curves and *D_av_* values for the multiple probes tested. The lipid-anchored probes are driven by their intrinsic partitioning preferences: PM-EGFP and Lyn-EGFP shift to slower diffusion in modulated Lo-like domains, whereas EGFP-GG and S15-EGFP shift to faster diffusion in proximal Ld-like regions (Figure 3B; summarized together with other probes in Figure 7A). Stabilization of an Lo-like environment that encompasses the clustered FcεRI may represent a thermodynamic adjustment, such as overcoming a hydrophobic mismatch of the collected TMDs and surrounding lipids (65). This adjustment may be accomplished by dynamic recruitment of the saturated lipids from the Ld-like regions, thereby establishing a more distinctive, phase-like membrane organization in the Ag-stimulated steady-state (18, 19, 66).

**Figure 7.**
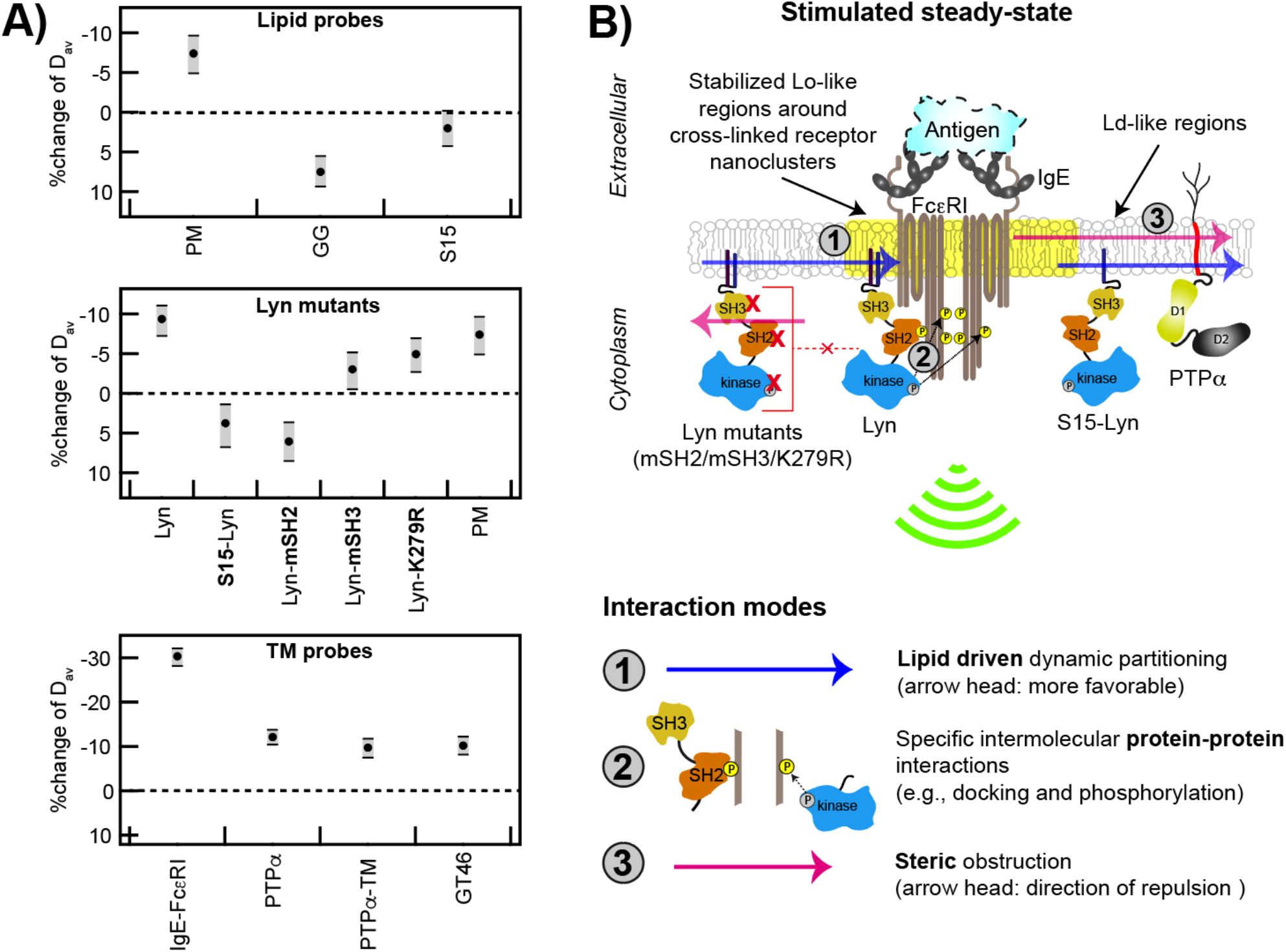
Ag-crosslinking of IgE-FcεRI stabilizes surrounding Lo-like nanodomains, causing dynamic lipid-based and protein-based interactions that shift diffusion properties of signaling components and lead to supra-threshold phosphorylation by Lyn. A) Stimulated changes in *D*_av_ for passive lipid probes, Lyn variants, IgE-FcεRI, PTPα, and TM probes. B) Proposed interaction modes leading to functional coupling of Lyn with clustered FcεRI: Stabilized Lo-like environment preferentially includes Lo-preferring Lyn and excludes Ld-preferring S15-Lyn and PTPα (interaction mode 1). Preferentially proximal Lyn (interaction 1) phosphorylates clustered FcεRI via its kinase module and then binds to pTyr via its SH2 module as facilitated by its SH3 module (interaction mode 2); these cumulative interactions stabilize the coupling. Lyn variants with impaired kinase, SH2 or SH3 modules are sterically hindered by cytoplasmic segments of clustered FcεRI (interaction mode 3). PTPα preferentially excluded from Lo-like environments (interaction mode 1) is further limited in access to FcεRI-pTyr because of steric obstruction by clustered FcεRI-TMDs (interaction mode 3).

This remodeling of membrane organization would facilitate lipid-driven sorting of Lo-preferring (e.g., Lyn) from Ld-preferring (e.g., PTPα) components as previously proposed (64). However, quantitative comparison of CDF curve shifts for Lyn-EGFP and PM-EGFP indicates that Lyn’s cytosolic protein modules (SH2, SH3, kinase) are also involved. As described previously (33), the fit parameters extracted from the *D* CDFs assess regions where a given probe interacts with other membrane constituents more strongly (interaction-rich Px units, characterized by *D*_slow_) and more weakly (interaction-poor Px units, characterized by *D*_fast_) (Figure 2C). Stimulated changes of these two parameters and their relative fraction (*F_slow_, F_fast_* = (1-*F_slow_*)) (Figure S3) provide further insights on modulation of the membrane environment as experienced by a particular probe. Notably, both *D*_fast_ and *D*_slow_ decrease for Lyn-EGFP while only *D*_slow_ decreases for PM-EGFP after antigen stimulation. A reasonable interpretation is that IgE-FcεRI clusters in the interaction-poor Px units do not adequately stabilize the Lo-like nanodomains to have an impact on the diffusion of PM-EGFP, which undergoes only lipid-based interactions. However, even relatively weak stabilization of Lo-like nanodomains in the interaction-poor Px units is sufficient to decrease the diffusion of Lyn-EGFP in these units due to its additional protein-based interactions with clustered IgE-FcεRI. ImFCS measurements of structural variants of Lyn and PTPα reveal a more complicated mechanism as depicted in Figure 7B and discussed further below.

### Lyn access to Ag-crosslinked IgE-FcεRI requires lipid-based filtering followed by protein binding

Comparing Lyn-EGFP to S15-Lyn-EGFP demonstrates that lipid-based sorting into stabilized Lo-like domains is the primary requirement for Lyn’s capacity to couple functionally with IgE-FcεRI. Unlike Lyn-EGFP and PM-EGFP, *D_av_* values and CDF curves for S15-Lyn-EGFP shift to faster diffusion after Ag addition, similarly to EGFP-GG and S15-EGFP (Figures 4 and 7A). This comparison shows that interactions mediated by Lyn’s cytosolic protein modules do not serve to slow diffusion unless Lyn’s PM anchor steers it into stabilized Lo domains. We infer that the slowed diffusion of Lyn-EGFP corresponds to Lyn partitioning into the Lo-like domains that surround Ag-crosslinked IgE-FcεRI (Figure 7B, interaction mode 1 (15)), followed by phosphorylating tyrosines in cytosolic segments of FcεRI (pTyr), and transiently binding to these pTyr via its SH2 module (Figure 7B, interaction mode 2; (67)). In comparison, S15-Lyn does not undergo interaction mode 1 and PM does not undergo interaction mode 2 (Figure 7B). These interactions differences, manifest in contrasting diffusion shifts after Ag addition (Figure 7A), are also manifest functionally: Ag addition triggers tyrosine phosphorylation mediated by Lyn-EGFP but not by S15-Lyn-EGFP (Figure 4D).

ImFCS measurements on Lyn probes with variations in cytosolic protein modules further show that protein interactions are involved in appropriate coupling with Ag-crosslinked IgE-FcεRI. We found that if SH2-mediated binding to phosphorylated FcεRI is prevented by a mutation in this module (53), then the diffusion of this Lyn-mSH2-EGFP shifts faster (rather than slower) after Ag addition (Figure 7A). These results suggest that the clustered cytosolic segments of Ag-crosslinked FcεRI sterically exclude the cytosolic protein modules of this Lyn variant (Figure 7B, interaction mode 3) to counteract lipid-based partitioning of its PM anchor. This interpretation is consistent with results with other Lyn variants. Unlike Lyn-mSH2-EGFP, the *D* CDFs and *D_av_* values for Lyn-K279R-EGFP and Lyn-mSH3-EGFP shift to slightly slower diffusion after Ag-crosslinking of IgE-FcεRI, but the extent of this negative shift is less than that for Lyn-EGFP and PM-EGFP (Figures 4 and 7A). We expect kinase-inactive Lyn-K279R-EGFP to be steered to stabilized Lo-domains (via PM anchor; interaction mode 1) resulting in slower diffusion, and this variant has the capacity to bind to FcεRI tyrosines that are phosphorylated by endogenous Lyn in these cells (interaction mode 2). However, endogenous Lyn is likely to be more competitive for proximal binding to the tyrosines it phosphorylates, and accordingly, Lyn-K279R-EGFP is probably more sensitive to steric exclusion by the clustered cytosolic segments of FcεRI (interaction mode 3).

The Lyn-mSH3-EGFP variant has a PM anchor, kinase activity and an intact SH2 module. However, the cytosolic SH3 module, which connects the PM anchor to SH2 and kinase modules, has been found to provide conformational plasticity of Lyn cytosolic segments for optimal catalytic activity and subsequent binding to pTyr in FcεRI (68). The impaired SH3 domain in Lyn-mSH3-EGFP is expected to limit this optimizing effect rendering this variant more susceptible to steric exclusion by the clustered FcεRI. These ImFCS results are consistent with our previous observations that Lyn-mSH2-EGFP is not recruited and Lyn-mSH3-EGFP only weakly recruited to micron-scale IgE-FcεRI clusters that form when cells are placed on antigen-micropatterned surfaces (53). In contrast, both Lyn-EGFP and PM-EGFP are recruited to these micropatterns (17, 53). Overall, the balance among interaction modes 1, 2, and 3 after Ag addition results in slower diffusion for Lyn-K279R-EGFP and Lyn-mSH3-EGFP but distinctively smaller shifts than for Lyn-EGFP (Figure 7A and B). We conclude that after Ag crosslinking of IgE-FcεRI, the primary coupling interaction is Lo-preference of Lyn’s PM anchor to facilitate its phosphorylation of FcεRI cytosolic segments followed by binding of its SH2 module to stabilize further the interaction. Despite these stabilizing effects the interactions are dynamic and relatively weak, such that the overall slowing of Lyn-EGFP diffusion is subtle (10% reduction in *D_av_*; Figure 7A).

### PTPα access to Ag-crosslinked IgE-FcεRI is reduced by lipid-based filtering and further limited by steric exclusion

As expected from previous results (24), PTPα-EGFP prefers an Ld-like environment as reflected by its detergent solubility (low *R* value; Figure 6A) relative to Lyn-EGFP (Figure 5A). It is surprising that the *D* CDF curves (Figure 6B,C) and *D_av_* (Figure 7A) show shifts to slower diffusion after Ag addition even though net localization of PTPα-EGFP in Ld-like regions, away from Lo-like domains stabilized around the clustered FcεRI (Figure 7B, interaction mode 1). Our ImFCS measurements further indicate that PTPα is sterically hindered from diffusing through the clustered TM segments of FcεRI (Figure 7B, interaction mode 3), unlike lipid-anchored PM-EGFP. That 70% of Ag-crosslinked IgE-FcεRI are immobilized would enhance this effect (Figure 1C) (60). The dominance of FcεRI-TMDs in excluding PTPα is indicated by similarly slowed diffusion of a passive TM probe (YFP-GL-GT46) and a PTPα variant (PTPα-E-TM-EGFP) without cytosolic protein modules (Figure 7A). These ImFCS results point to the importance of both lipid-based and steric exclusion processes in protecting Ag-crosslinked IgE-FcεRI, phosphorylated by Lyn, from dephosphorylation from a TM phosphatase. We cannot rule out other causes for the stimulated diffusional shifts of TM probes we evaluated. However, similar steric hindrance of TM phosphatase and its impact on B cell signaling was recently reported (43).

## CONCLUSION

As depicted by interaction modes in Figure 7B, a coordinated synergy of lipid-based and protein-based interactions explains how Ag-crosslinking of IgE-FcεRI leads to its supra-threshold tyrosine phosphorylation, both by facilitating access by Lyn kinase and limiting access by a TM phosphatase. Based on many studies in our and other laboratories, we take the view that the resting cell is poised to respond a specific stimulus and the change in membrane organization to initiate signaling is subtle (15). The subtlety of the change has made detection challenging, requiring super-resolution imaging, SPT, and other technically difficult approaches. The strength of the mechanism proposed in Figure 7B rests on precise ImFCS measurements of small but distinctive diffusion shifts stimulated by Ag for multiple structural variants of the key signaling components and passive lipid probes. Although this suggested mechanism is based primarily on diffusion measurements, these are both internally consistent and consistent with previous studies cited here. In particular, we showed directly and provided the strongest evidence to date that Lyn’s lipid-based steering is necessary to initiate tyrosine phosphorylation. Overall, we demonstrated the relative ease of applying ImFCS, using multiple probes and conventional fluorophores, to dissect contributions of structural features to weak interactions that collectively have decisive impact in stimulated TM signaling. We expect that ImFCS and the experimental strategies described herein will be widely applicable to advance understanding of TM signaling where plasma membrane organization is likely to play an integral role.

## ACKNOWLEDGEMENTS

We thank Alex Batrouni and Prof. Jeremy Baskin (Cornell University) for the access of their confocal microscope for FRAP measurements. We thank Henry Phan and Boyu Yin for discussions on the DRM preparations. This work is supported by National Institute of General Medical Sciences Grant R01GM117552. The content is solely the responsibility of the authors and does not necessarily represent the official views of NIGMS or NIH.

## Notes

### Competing Interest Statement

The authors have declared no competing interest.

