## Supporting Information for "Lipid-based, protein-based, and steric interactions synergize to facilitate transmembrane signaling stimulated by antigen-clustering of IgE receptors"

#### MATERIALS AND METHODS

##### Reagents

Minimum essential medium (MEM), F-12 medium, Opti-MEM, Trypsin-EDTA (0.01%) and gentamicin sulfate were obtained from Life Technologies (Carlsbad, CA). Fetal Bovine Serum (FBS) was purchased from Atlanta Biologicals (Atlanta, GA). Anti-phosphotyrosine antibody clone 4G10 was purchased from Millipore (Billerica, MA). Alexa Flour 633 (AF633) anti-mouse immunoglobulin G 2b (IgG2b) secondary antibody was purchased from Life Technologies (Carlsbad, CA). Phosphate Buffered Saline (PBS) was obtained from Sigma Aldrich. Gentamicin and Geneticin (G418 sulfate) were purchased from Thermo Fisher Scientific (Waltham, MA). Alexa Fluor 488 (AF488) NHS ester (Invitrogen) was used to fluorescently label monoclonal anti-DNP (2,4-dinitrophenyl) immunoglobulin E (IgE) yielding AF488-IgE as described previously (1). The antigenic multivalent ligand, DNP-BSA, was prepared by conjugating DNP sulfonate (Sigma-Aldrich) to bovine serum albumin (BSA) (2). Phorbol 12,13-dibutyrate (PDB) was obtained from Sigma-Aldrich (St. Louis, MO). Stock solution of PDB was prepared in DMSO and stored at -80°C.

##### Plasmids

The new DNA constructs created during this study are described below.

<u>PTPα-EGFP and PTPα-mCherry</u>

The initial PTPα-mEos3.2 was created by PCR using the PTPα-HA plasmid, provided by David Shalloway (Cornell University) and primers (forward sequence) 5’-CGCCGCTAGCGGCCACCATGGATTCC-3’ and (reverse sequence) 5’-TGTCCTCGAGCTTGAAGTTGGCATAAT-3’. The fragment was ligated into the mEos3.2-N1 vector using generated 5’-NheI and 3’-XhoI restriction sites.

PTPα-mCherry was generated by exchanging the fluorescence protein in the PTPα-mEos3.2 construct with the mCherry sequence by digestion with 5’-XhoI and 3’-NotI.

PTPα-EGFP was constructed by exchanging the fluorescence protein in the PTPα-mCherry construct with the EGFP sequence by digestion with 5’-XhoI and 3’-NotI.

<u>PTPα-E-TM-EGFP</u>

This construct is created by deleting the intracellular part of the PTPα-EGFP construct. The extracellular and transmembrane portions of PTPα with 5’-NheI and 3’-XhoI sites was created by PCR with primers (forward sequence) 5'-AAAAAGCTAGCGGCCACCATGGATTCCTGG-3' and (reverse sequence) 5'-AAAAACTCGAGTCTGGCCAGAAGTGGCACACTCTGG-3'. The fragment was ligated in to pEGFP-N1 (Clontech Laboratories, Palo Alto, CA).

S15-Lyn-EGFP

The S15-Lyn-EGFP construct, where the first 15 amino acids of Lyn are replaced by the first 15 amino acids of Src-kinase, was generated in two consecutive cloning steps. In a first step a truncated version of the wt-Lyn was created by PCR with primers (forward sequence) 5'-AAAACTGCAGGGAGTAGATATGAAGACTCAACCAGTTCCTGAATC-3' and (reverse sequence) 5'-AAAAAGGATCCGCCGGTTGCTGCTG-3' and subcloned into the pEGFP-N1 vector via the 5'-PstI and 3'-BamHI sites.

In a second step the Src-kinase N-terminal portion was inserted into the above generated plasmid by using 5'-XhoI and 3'-PstI and annealed complementary oligos 5'-TCGAGATGGGGAGCAGCAAGAGCAAGCCCAAGGACCCCAGCCAGCGCCGGCTGCA-3' and 5'-GCCGGCGCTGGCTGGGGTCCTTGGGCTTGCTCTTGCTGCTCCCCATC-3'.

Lyn-K279R-EGFP

The Lys to Arg (at position 279 in the kinase domain of Lyn) Lyn mutant (Lyn-K279R-EGFP) of wt Lyn-EGFP (3) was generated performing site-directed mutagenesis with primers (forward sequence) 5'-GCACAAAAGTGGCTGTAAGGACCCTCAAGCCTGG-3' and (reverse sequence) 5'-CCAGGCTTGAGGGTCCTTACAGCCACTTTTGTGC-3'.

**RBL cell culture, transfection, sensitization and stimulation**

RBL-2H3 mast cells (for brevity, RBL cells) were cultured in growth medium (80% MEM supplemented with 20% FBS and 10 mg/L gentamicin sulfate) at 37°C and 5% (v/v) $CO_2$ environment.

*Chemical transfection:* RBL Cells in a confluent 25 $cm^2$ flask were washed once with 2 mL Trypsin-EDTA, detached with 2 mL Trypsin-EDTA for 5 min at 37°C and 5% (v/v) $CO_2$ environment. The Trypsin-EDTA is quenched with 8 mL of growth medium (~$10^6$ cells/mL). About 20,000 cells were homogeneously spread in a 35 mm MatTek dish (Ashland, MA) containing 2 mL growth medium and allowed to grow overnight. MatTek dishes containing the adherent cells were transfected using FuGENE HD transfection kit (Promega). For one MatTek dish, plasmid DNA (0.5 – 1 µg) and FuGENE (3 µL FuGENE/µg DNA) were first mixed in 100 µL Opti-MEM medium and incubated at room temperature for 15 min. Next, MatTek dishes containing cells were washed once and covered with 1 mL Opti-MEM. The DNA/FuGENE complex was spread evenly over the cells and incubated for 1 hr, followed by incubation with pre-warmed PDB (1 mL, 0.1 µg/mL) for 3 hr at 37°C in 5% (v/v) $CO_2$ environment. Finally, 2 mL of growth medium was added to each MatTek dish after discarding Opti-MEM. The transfected cells were cultured for 18 – 22 hr at 37°C in 5% (v/v) $CO_2$ environment before DRM preparation or live cell imaging or FRAP measurements. The chemically transfected plasmids used in this study encode the following proteins: PM-EGFP (4), EGFP-GG (4), S15-EGFP (5), YFP-GL-GPI

(6), YFP-GL-GT46 (6), Lyn-EGFP (3), Lyn-mSH2-EGFP (7), Lyn-mSH3-EGFP (7), Lyn-K279R-EGFP (8), and S15-Lyn-EGFP.

Lyn-mSH2-EGFP, Lyn-mSH3-EGFP, and Lyn-K279R-EGFP are point mutants of Lyn-EGFP to disable functions of SH2, SH3, and kinase modules respectively. The mutation sites are: Arg to Ala at position 135 (Lyn-mSH2-EGFP), Try to Ala at position 78 (Lyn-mSH3-EGFP), and Lys to Arg at position 279 (Lyn-K279R-EGFP).

PTPα-E-TM-EGFP was transfected using lipofectamine 2000 reagent kit (Thermo Fisher Scientific (Waltham, MA)) following manufacturer's protocol. For one MatTek dish prepared as before (for the FuGENE-based transfection), 1 µg plasmid and 4 µL lipofectamine reagent was used.

*Electroporation:* RBL cells in a confluent 75 $cm^2$ flask were washed and trypsinized for 8 min at 37°C and 5% (v/v) $CO_2$ environment with 3 mL Trypsin-EDTA. The detached cells were resuspended in 7 mL of growth medium and centrifuged to remove the medium. The cell palette ($15\times10^6$ cells) was resuspended in 1.5 mL of cold electroporation buffer (137 mM NaCl, 2.7 mM KCl, 1.0 mM $MgCl_2$, 1 mg/ml glucose, and 20 mM HEPES; pH 7.4). Next, 10 µg of plasmid DNA was thoroughly mixed with 500 µL of the resuspended cells in an electroporation cuvette (Bio-Rad). This cuvette was subject to an electroporation pulse (280 V, 950 µF) using a Gene Pulser X (Bio-Rad) electroporation module. The electroporated cells were then added to 6 mL of growth medium, mixed thoroughly, and deposited in MatTek dishes (2 mL/dish). The cells were allowed to attach on the dish for 3 hr at 37°C and 5% (v/v) $CO_2$ environment following which the medium was replaced with fresh growth medium. The cells were cultured for 24 hr to recover before proceeding to the next sample preparation steps. The electroporated plasmids used in this study encode the following proteins: PTPα-EGFP.

*Cell sensitization and stimulation:* RBL cells were washed twice with Buffered Salt Solution (BSS: 135 mM NaCl, 5.0 mM KCl, 1.8 mM $CaCl_2$, 1.0 mM $MgCl_2$, 5.6 mM glucose, and 20 mM HEPES; pH 7.4) and sensitized with 2 µg/mL of anti-DNP IgE (for transfected cells to monitor stimulation-induced changes of the transfected probe) or a mixture of 0.5 µg/mL AF488-IgE and 1.5 µg/mL of anti-DNP IgE (for untransfected cells to test the stimulation-induced changes of FcεRI) prepared in BSS for 40 min at room temperature. The cells were washed twice with BSS and stimulated with 0.5 µg/mL DNP-BSA antigen (Ag) for 15 min at room temperature. Finally, the cells were washed twice with BSS and imaged in fresh BSS or subjected to detergent resistant membrane (DRM) preparation.

**Immunostaining of Chinese Hamster Ovary (CHO) cells stably transfected with FcεRI (CHO-FcεRI) and imaging:**

CHO-FcεRI cells (9) were maintained in 80% F-12 and 20% FBS medium containing 50 mg/mL Geneticin (G418 sulfate) and 1 mg/mL Gentamycin antibiotics in 37°C and 5% $CO_2$ environment. The expression of FcεRI in these cells were routinely monitored by labelling them with AF488-IgE.

For immunostaining, CHO-FcεRI cells were grown to 70-80% confluency in MatTek dishes (Ashland, MA) and were transfected using Mirus TransIT-2020 (Mirus Bio, Madison, WI) reagent kit following manufacturer's protocol. Typically, 1 μg of Lyn-EGFP or S15-Lyn-EGFP plasmid along with 2 μg PTPα-mCherry plasmid and 6 μL of Mirus reagent was used per MatTek dish. The cells were incubated ~22-24 hours with the plasmid/Mirus mixture in 37°C and 5% $CO_2$ incubator. The cells were then washed twice with BSS and sensitized with 2 μg/mL IgE in BSS for 40 minutes at room temperature. The excess IgE was washed with BSS, and then the cells were incubated with either fresh BSS (resting condition) or 0.9 μg/mL DNP-BSA antigen (stimulated condition) for 5 minutes at 37°C (10). Following this, the cells were washed once with BSS and twice with PBS buffer and fixed with 4% paraformaldehyde and 0.1% glutaraldehyde in PBS for 10 minutes at room temperature. The fix was quenched with blocking buffer 10 mg/mL bovine serum albumin (BSA) in PBS. The fixed cells were then permeabilized and labelled with anti-phosphotyrosine antibody (4G10) solution (5 μg/mL 4G10, 0.1% Triton X-100, and 10 mg/mL BSA in PBS) for 1 hour at room temperature. The dishes were then washed multiple times with blocking buffer (10 mg/mL BSA in PBS) followed by incubation with secondary antibody (1 μg/mL Alexa Flour 633 (AF633) anti-mouse immunoglobulin G 2b (IgG2b) antibody, 0.1% Triton X-100, and 10 mg/ml BSA in PBS) for 1 hour at room temperature. The dishes were washed multiple times with blocking buffer and stored in PBS at 4°C until imaging.

Fluorescence imaging of the cells was performed using the epi-fluorescence microscope (for TIRF imaging) described below. We used 641 nm laser (Coherent, Santa Clara, CA) and PLAN, 10×, 0.22 NA objective to excite the sample. The fluorescence images were recorded by an electron multiplying charge coupled device (EMCCD) camera (black illuminated Andor iXON3 897, pixel size 16 μm, Andor Technology, Belfast, UK) after being filtered by a ZET405/488/561/640m emission filter (Chroma technology). Generally, more than 100 cells from 7-10 fields of view (FOVs) were imaged per dish. Average fluorescence of individual cells was determined after background correction using FIJI (11). Average fluorescence of multiple regions of interest outside cells in a given FOV was used as background. For each pair of samples (resting and stimulated conditions), at least three independent experiments were performed. The fold change of phosphorylation (as quantified from the fluorescence of the AF633 labelled secondary antibody against 4G10) of stimulated cells relative to the resting cells were quantified from each biological replica.

**Fluorescence Recovery after Photobleaching (FRAP)**

FRAP experiments were performed in Zeiss 710 confocal microscope equipped with a high power 488 nm laser source, an oil-immersion, 40×, 1.2 NA objective, and a sensitive photomultiplier tube detector. In a typical FRAP experiment, a region of interest (ROI) of 3.5×3.5 $\mu m^2$ (bleached ROI) on the ventral surface of fluorescently labelled cell was photobleached with high power 400 nm laser (100% laser power). The fluorescence recovery of this spot is recorded at low laser power (0.2% laser power). We also simultaneously recorded fluorescence of an unbleached spot of same size on the cell (reference ROI) and a spot outside the cell as background ROI. In addition, five time-lapse images of all three ROIs were taken before photobleaching of the bleached ROI to create normalized FRAP curves. All measurements were carried out at room temperature.

The experimental fluorescence counts against time of the bleached ROI is background-corrected (by subtracting the background ROI counts) and normalized using the fluorescence counts of the reference ROI and pre-bleaching intensity counts such that normalized fluorescence before photobleaching equals to 1 ($F_{normalized}(t < 0) = 1$; pre-bleaching) and at the time of photobleaching is zero ($F_{normalized}(t = 0) = 0$; at the bleaching). This is done using FRAPanalyser (http://actinsim.uni.lu/eng/Downloads/FRAPAnalyser) (12). The normalized recovery curve (normalized intensity ($F_{normalized}(t \geq 0)$) against time ($t \geq 0$ )) post-bleaching was fitted with a single-exponential model (Eqn S1) using Igor Pro (Version 8; WaveMetrics, OR, USA). The saturation value of the fitted curve at long time (i.e., when recovery is completed), $F_{max}$, is the mobile fraction while time scale of diffusion is given by the recovery time ($\tau_{1/2}$) of the bleached spot.

$$F_{normalized}(t \geq 0) = F_{max}\left[1 - \exp\left(-\frac{t}{\tau_{1/2}}\right)\right] \qquad \text{(S1)}$$

FRAP experiments are performed on multiple cells for a given condition from at least three independent samples. Recovery time and mobile fraction of individual cells were determined using Eqn S1. Statistical significance of these parameters between two conditions were done by Mann-Whitney test.

**Preparation of detergent resistant membrane (DRM) imaging samples**

The entire DRM preparation was done in an ice bath. First, a pair of MatTek dishes containing fluorescently labelled cells were first placed in the ice bath for 10 min followed by washing with BSS once. In the experiment dish (+TX100), the cells were treated with 1 mL of 0.04% (v/v) cold TX100 in BSS while the control dish (-TX100) was treated with 1 mL of cold BSS for 10 min. The cells were then fixed with 4% paraformaldehyde and 0.1% glutaraldehyde in PBS for 10 min followed by quenching by 10 mg/mL BSA in PBS for another 20 min. The fixed cells were washed with PBS and stored in fresh PBS at 4°C and imaged within 2 days.

**TIRF imaging**

Fluorescently labeled RBL cells were imaged with a home-built total internal reflection fluorescence microscope (TIRFM) (DMIRB, Leica Microsystems, Germany) equipped with an oil immersion objective (PlanApo, 100×, NA 1.47; Leica Microsystems, Germany), a 488 nm excitation laser (Coherent, Santa Clara, CA), and an electron multiplying charge coupled device (EMCCD) camera (black illuminated Andor iXON3 897, pixel size 16 µm, Andor Technology, Belfast, UK). The excitation laser beam was introduced and focused on the back focal plane of the objective by a pair of tilting mirrors and a dichroic mirror (ZT405/488/561/640rpc, Chroma Technology). The same set of mirrors was used to adjust the TIRF angle of the excitation beam to illuminate the ventral membrane. The fluorescence signal from the sample was recorded by the EMCCD camera after it passes through the same objective and the dichroic mirror and reflected to the camera chip after being filtered by an emission filter (ZET488/561m, Chroma

Technology). For both DRM and live cell imaging, 100 TIRF images were taken with 10 ms exposure time. Andor Solis software was used for the image acquisition. The laser power was 50 μW before objective. All measurements were carried out at room temperature.

**Quantification of DRM fraction imaging**

About 30 cells expressing a given probe were imaged for each of the +TX100 and -TX100 samples using the above TIRF imaging protocol. The average background-corrected fluorescence count of each cell was calculated by subtracting background (from a region outside the cell) from the average fluorescence signal of the region inside that cell using FIJI/ImageJ (11). This yields a range of background-corrected fluorescence counts per cell before (-TX100 sample) and after (+TX100 sample) detergent treatment. The distributions were subjected to non-parametric Mann-Whitney test to check whether the fluorescence of the cells from both samples belong to the same distribution. The null hypothesis ($P > 0.05$) for this test was that a randomly selected fluorescent values both a +TX100 and a -TX100 sample belong to the same distribution. In this case ($P > 0.05$), we consider the probe is completely detergent-resistant under the experimental condition. If a probe is detergent-soluble, the Mann-Whitney test between -TX100 and +TX100 samples returns a P value < 0.05. The extent of detergent-resistance is then quantified as the Resistance factor (*R*), which is calculated as (Eqn S2):

$$R = \frac{\text{Median background-corrected fluorescence of } (+)\text{TX100 sample}}{\text{Median background-corrected fluorescence of } (-)\text{TX100 sample}} \qquad \text{(S2)}$$

An *R* value of 1 suggests complete resistance (i.e., $P > 0.05$ between -TX100 and +TX100 samples) while *R* equals to zero for complete solubility of a probe in 0.04% TX100. An intermediate value of *R* indicates partial detergent-resistance. Typically, DRM imaging for a given probe was performed twice for both unstimulated and stimulated conditions.

The mean and error of *R* value were determined by bootstrapping with 50% of the data. Briefly, we first randomly sub-sampled 50% of the cells (e.g., 15 out of 30 cells) from both -TX100 and +TX100 samples from a biological replica. An *R* value was then determined from the median values of background corrected fluorescence per cell from these sub-samples according to Eqn S2. These sub-sampling steps were repeated for 10000 times and subsequently 10000 *R* values were determined. This method was then repeated for all biological replicas to obtain a total of 10000*n *R* values where n = number of biological replicas. The arithmetic average of these 10000*n *R* values is reported in the main text, while the error (± values) represent standard deviation/√n.

**Data acquisition of ImFCS and ACF analysis to determine diffusion coefficient (*D*) values**

The data acquisition protocol for ImFCS and following autocorrelation function (ACF) analysis were described previously (13). Briefly, a stack of 80,000 images from a ROI (40×40 to 50×50 pixels with pixel size of 160 nm in the object plane) on the ventral plasma membrane was recorded at an acquisition speed of 3.5 ms/frame using the TIRF microscope and EMCCD camera described above and saved as .fits or .tif file. All measurements were carried out at room temperature.

This raw image stack was further processed by a FIJI (11) plug-in for ImFCS (Imaging_FCS 1.491; available at http://www.dbs.nus.edu.sg/lab/BFL/imfcs_image_j_plugin.html). Raw temporal autocorrelation function (ACF) ($G(\tau)$) were computed from each 2×2 binned pixels (Px unit; length = 320 nm) of an image stack and fitted with Eqn S3 (14). This yields a map of lateral diffusion coefficient (*D*) values.

$$G(\tau) = \frac{1}{N}\left(\frac{\mathrm{erf}\big(p(\tau)\big) + \frac{\left(e^{-(p(\tau))^2-1}\right)}{\sqrt{\pi}\,p(\tau)}}{\mathrm{erf}\left(\frac{a}{\omega_0}\right) + \frac{\omega_0}{a\sqrt{\pi}}\left(e^{-\frac{a^2}{\omega_0^2}} - 1\right)}\right)^2 + G_\infty; \quad p(\tau) = \frac{a}{\sqrt{4D\tau + \omega_0^2}} \qquad \text{(S3)}$$

In the above equation, $G(\tau)$ is the ACF as a function of lag time ($\tau$), *N* is the number of particles diffusing within a Px unit, *D* is the lateral diffusion coefficient in the Px unit, *a* is the length of the Px unit in the object plane (320 nm), $\omega_0$ is the point spread function (PSF) of the microscope, $G_\infty$ is the convergence value of $G(\tau)$ at very large lag times. We used *N, D* and $G_\infty$ as fit parameters, and $\omega_0$ was experimentally determined using the method described previously (15).

*Construction of cumulative distribution function (CDF) of D and determination of Stimulated %change of $D_{av}$ as shown in Figure 2-6 in the main text*

After combining *D* values obtained from multiple cells over multiple preparations for a given condition (red: – Ag or black: + Ag), this large data set was 30 times resampled by bootstrapping with 50% of the data (See Appendix in the end of the SI for more detail on the bootstrapping analysis). Individual CDFs were then created from each bootstrapped sub-sample, and these are overlaid in Figures 2-6 in the main text. The associated % change of $D_{av}$ was determined as follows: First, mean values of one, randomly selected, bootstrapped sub-sample for each of – Ag and + Ag conditions are determined ($D_{BS,-Ag}$ and $D_{BS,+Ag}$). The stimulated %change of $D_{av}$ for this pair is calculated as: $(D_{BS,+Ag} - D_{BS,-Ag})$*100%/$D_{BS,-Ag}$. This process for randomly selected pairs is repeated 10,000 times, and a histogram of %change of $D_{av}$ is created.

#### SI FIGURES

##### Figure S1

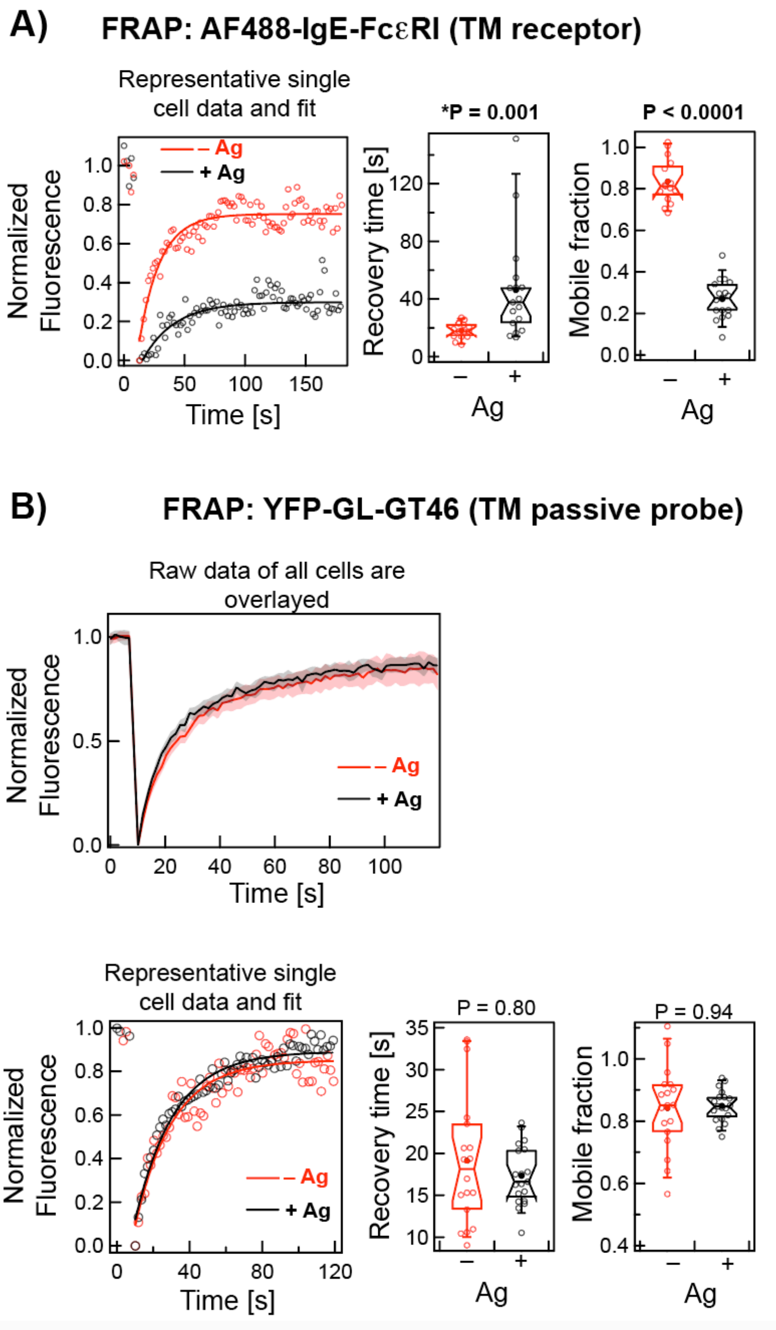


**Figure S1.** FRAP data for A) TM receptor AF488-IgE-FcεRI, and B) Ld-preferring TM probe, YFP-GL-GT46 in –/+ Ag conditions in RBL cells. The left panel in A and the bottom left panel of B show representative raw fluorescence recovery curves (circle) and corresponding fits (solid line using Eqn S1) of specified probe under – Ag (red) and + Ag (black) conditions. The middle and right panels show the fitted values of recovery time and mobile fraction from multiple cells, respectively, as box plots (Eqn S1 defines these parameters). The box height corresponds to 25$^{th}$ to 75$^{th}$ percentile and error bars represent 9$^{th}$ to 91$^{st}$ percentile of entire data set. Mean and median values are represented as solid circle and bar, respectively, located inside the box. The notches signify 95% confidence interval of the median. The top panel of B shows normalized FRAP curves of YFP-GL-GT46 from multiple cells in – Ag (pink) and + Ag (grey) conditions. The solid red and black curves are the average of the pink and grey curves respectively. Number of cells for AF488-IgE-FcεRI: 15 (– Ag) and 17 (+ Ag); and for YFP-GL-GT46: 18 (– Ag) and 19 (+ Ag).

#### Figure S2

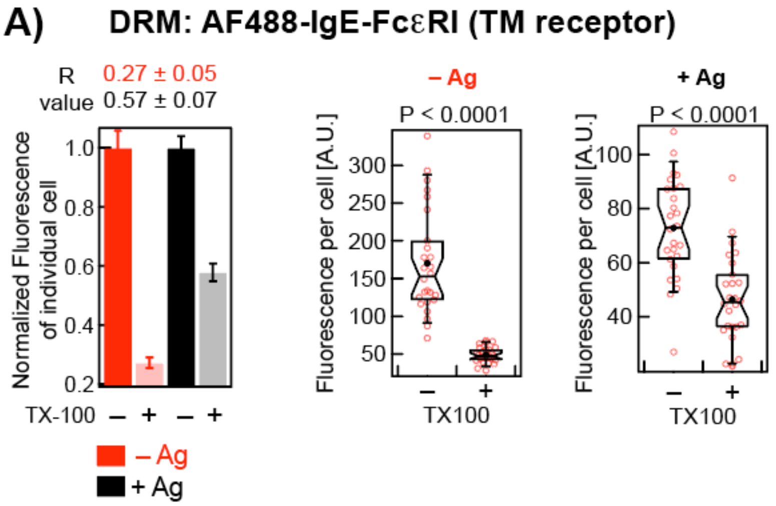


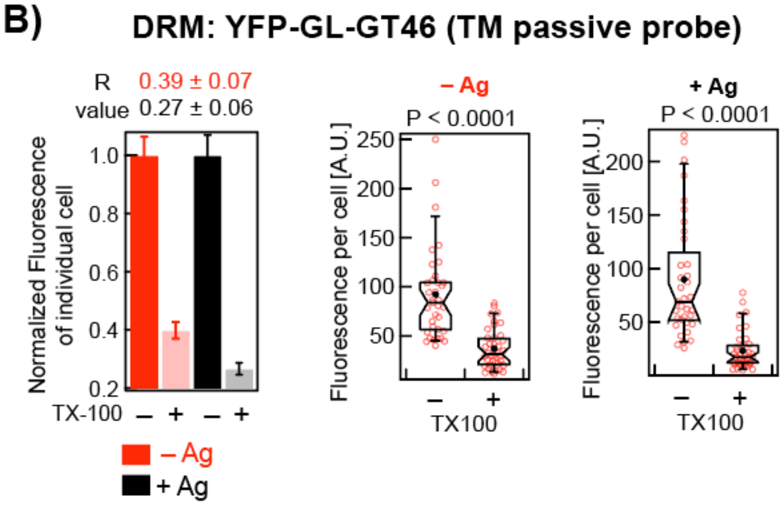


**Figure S2.** DRM results for A) TM receptor AF488-IgE-FcεRI, and B) Ld-preferring TM probe, YFP-GL-GT46 under –/+ Ag conditions in RBL cells. In the left panel, the relative loss of fluorescence and the corresponding *R* value (Eqn S2) after 0.04% TX100 treatment for the probes in –/+ Ag conditions for both probes. Each bar represents data from 60-90 cells from 2-3 independent experiments. The box plots in the right panel show raw fluorescence values of ~30 cells from a single representative experiment for each of –/+ Ag and –/+ TX100 conditions. Box parameters described in legend to Figure S1

**Figure S3**

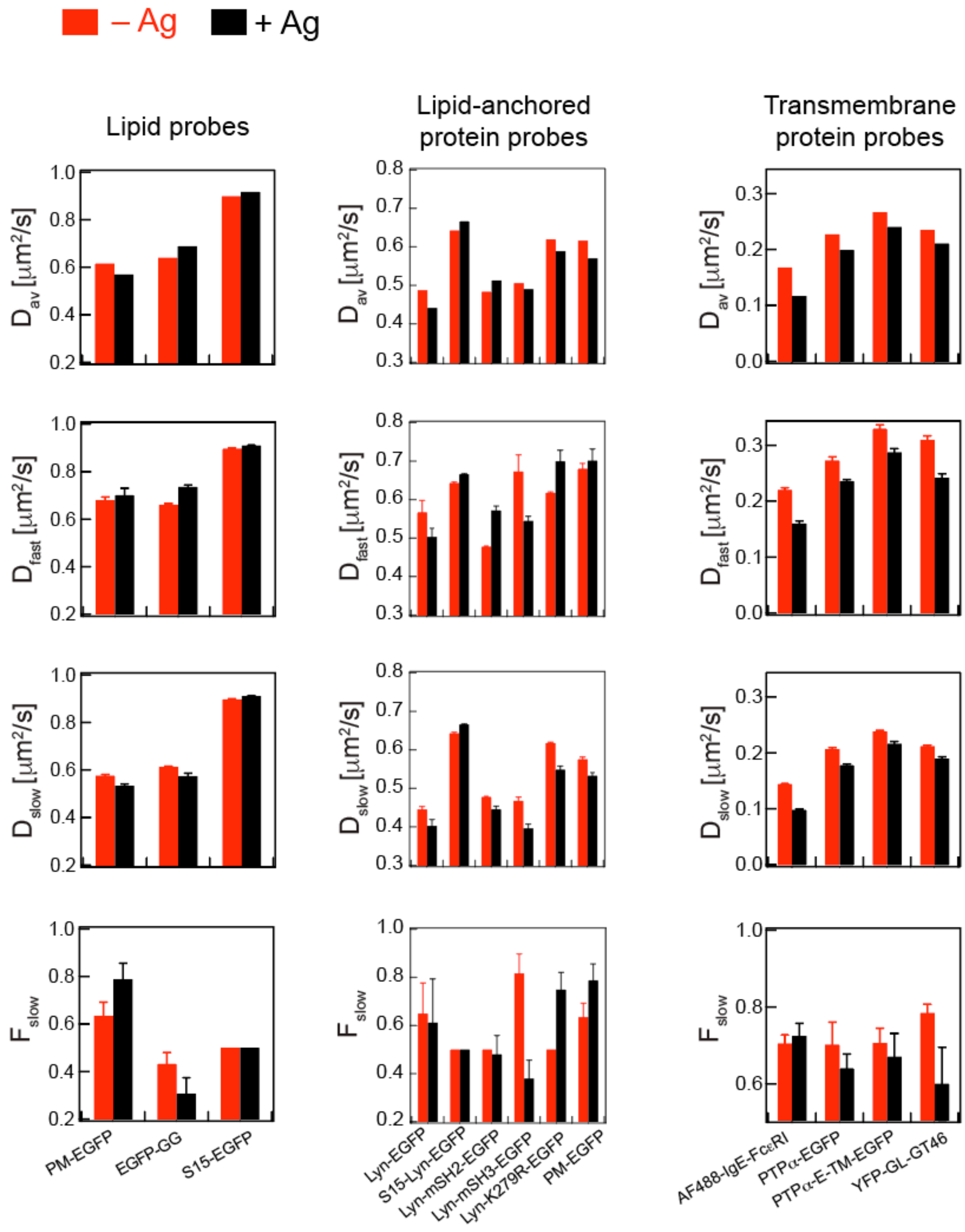


**Figure S3.** Fit results of bootstrapped CDFs obtained all probes tested in this study under –/+ Ag (red/black) conditions using Eqn A1 or A2 (SI Appendix). Error bars represent standard deviation from the fitting 30 bootstrapped CDFs individually.

**Figure S4**

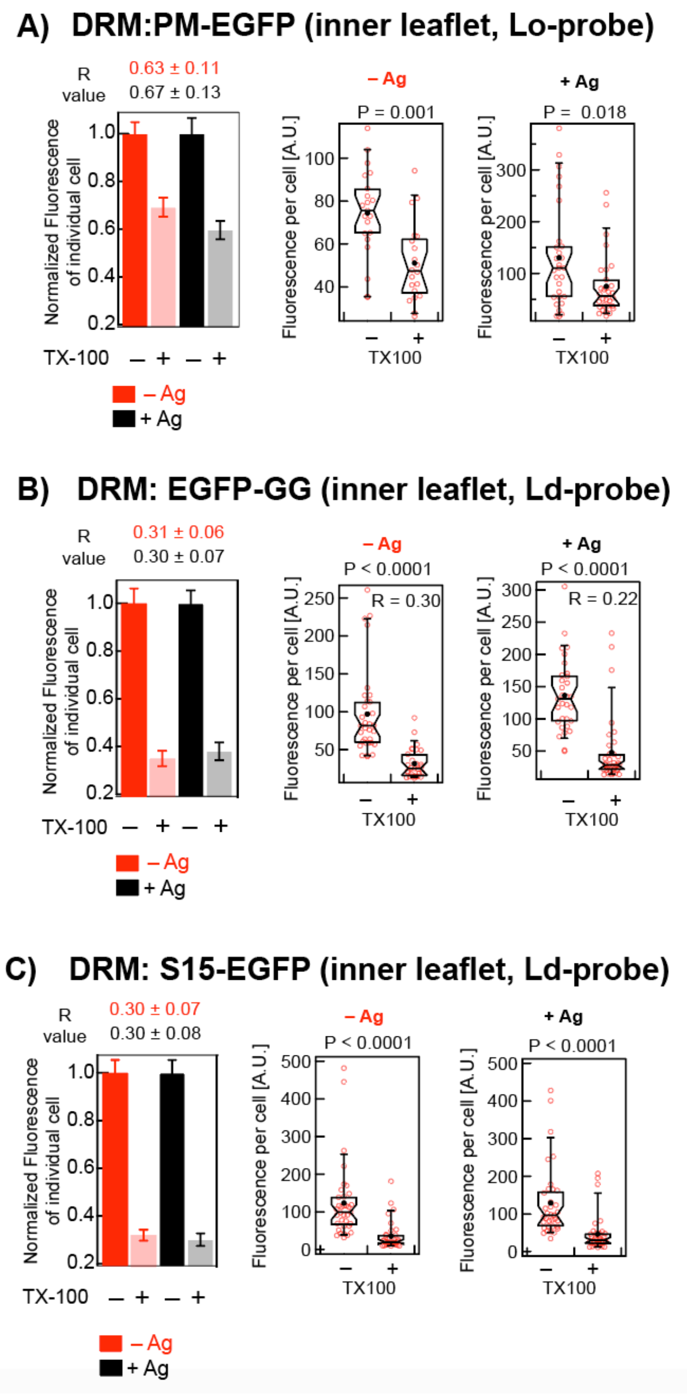


**Figure S4.** DRM results for the lipid probes under –/+ Ag conditions in RBL cells: A) PM-EGFP, B) EGFP-GG, and C) S15-EGFP. Left panel shows relative loss of fluorescence and the corresponding *R* value (Eqn S2) upon 0.04% TX100 treatment for the probes in –/+ Ag conditions. Each bar represents data from 60-90 cells from 2-3 independent sample preparations. Right panels show box plots of raw fluorescence values of ~30 cells for each of –/+ TX100 conditions from a representative experiment. Box parameters described in legend to Figure S1

**Figure S5**

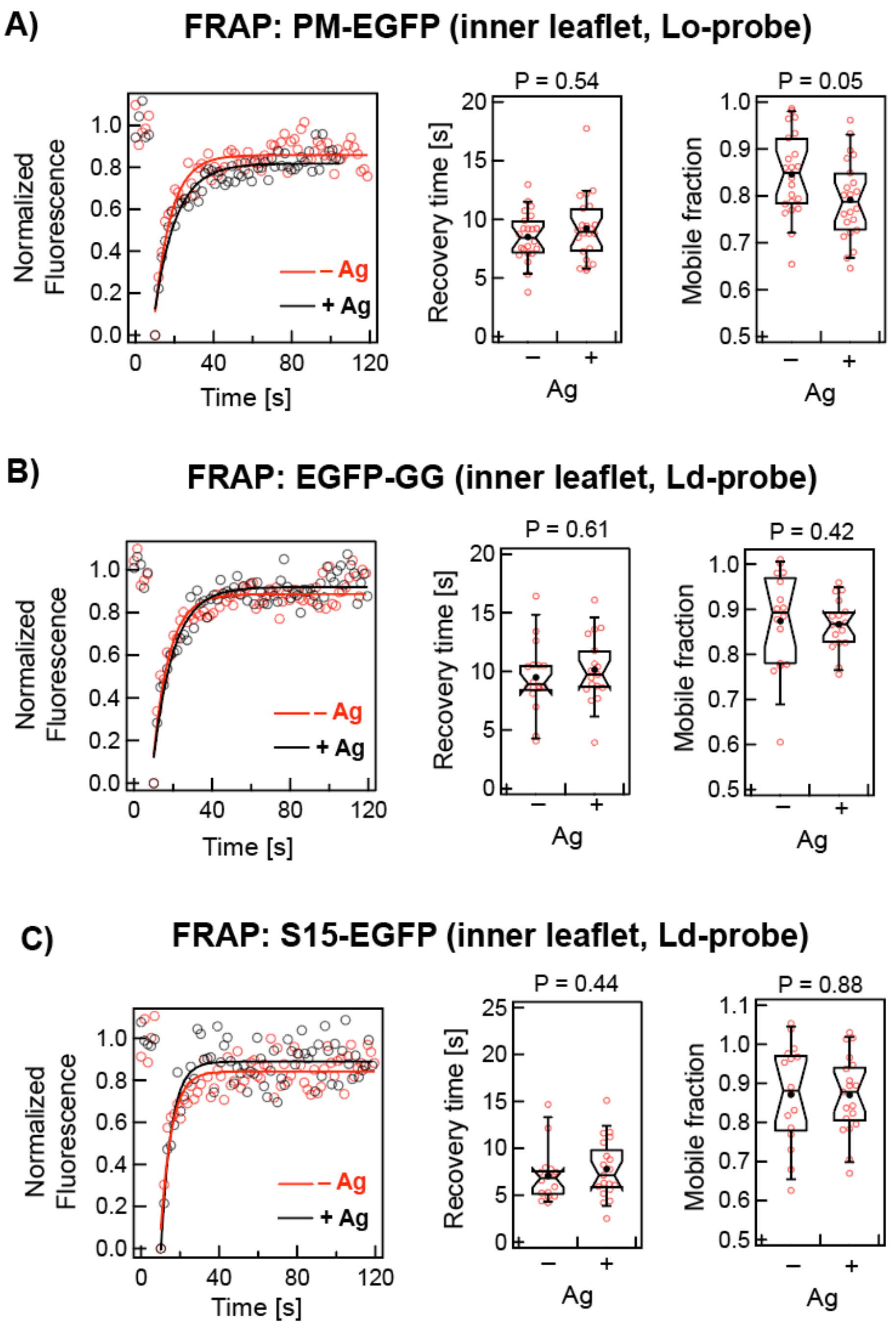


**Figure S5.** FRAP results for lipid probes under –/+ Ag conditions in RBL cells: A) PM-EGFP, B) EGFP-GG, and C) S15-EGFP. Left panels in A-C show representative raw fluorescence recovery curves (circle) and corresponding fits (solid line using Eqn S1) of the specified probe under – Ag (red) and + Ag (black) conditions. Right panels show fitted values of recovery time and mobile fraction, respectively, from multiple cells as box plots (Eqn S1 defines these parameters). Box parameters are described in legend to Figure S1. Number of cells for PM-EGFP: 22 (– Ag) and 22 (+ Ag); for EGFP-GG: 16 (– Ag) and 17 (+ Ag); for S15-EGFP: 16 (– Ag) and 19 (+ Ag).

**Figure S6**

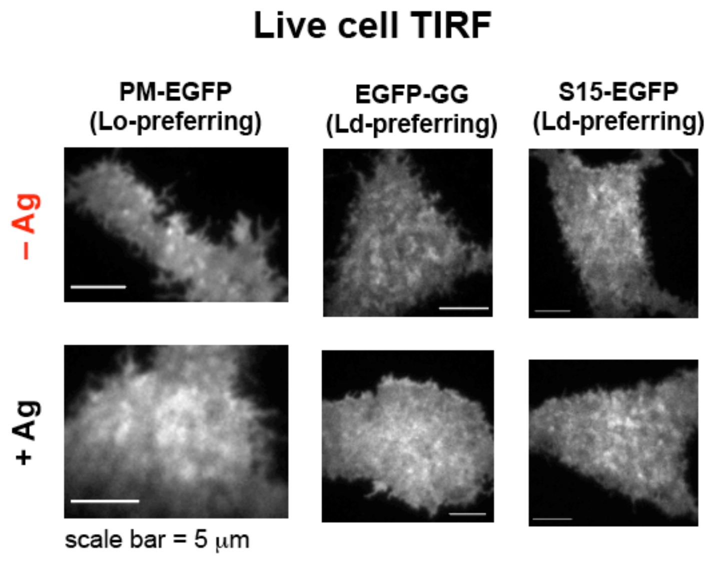


**Figure S6.** Representative live cell TIRFM images of specified lipid probes under –/+ Ag conditions in RBL cells.

### Figure S7

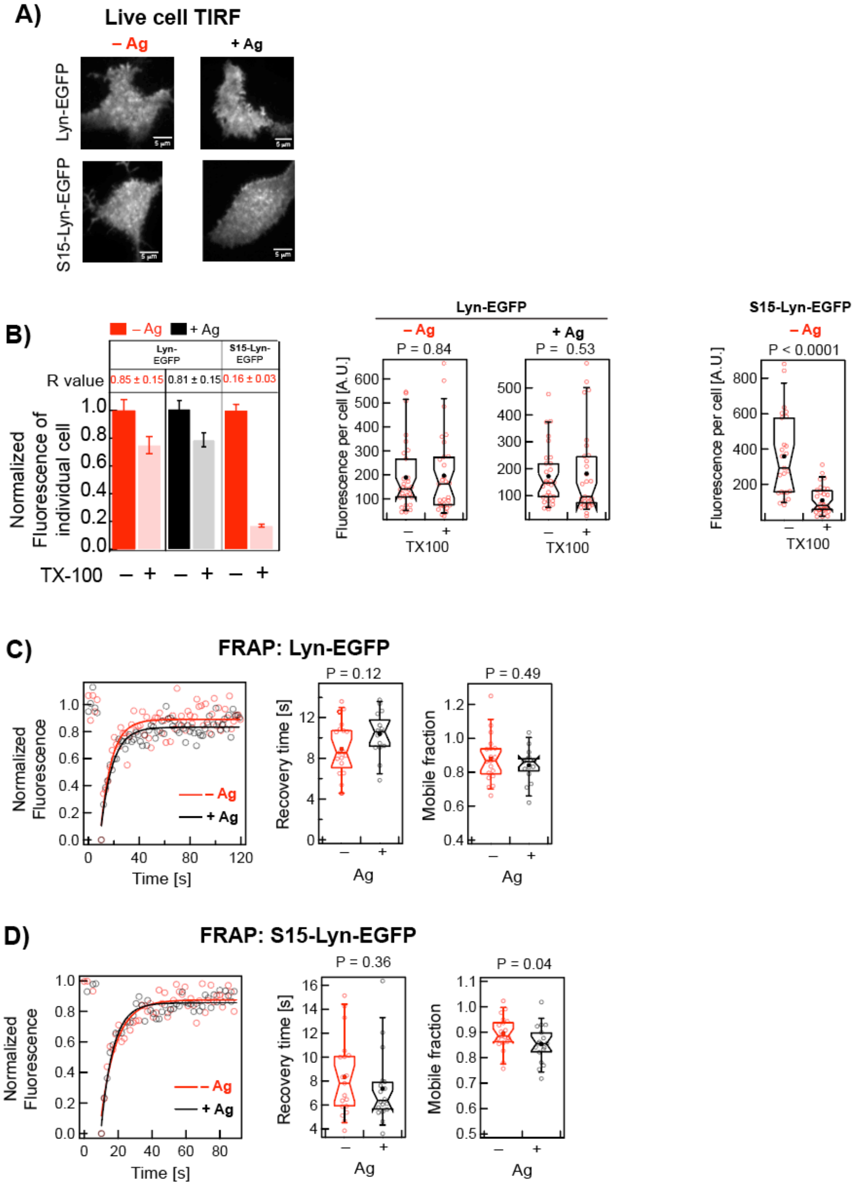


**Figure S7.** Comparison of the biophysical properties of Lyn-EGFP and S15-Lyn-EGFP under –/+ Ag conditions in RBL cells. A) Representative live cell TIRF images. B) DRM results: Left panels show relative loss of fluorescence and the corresponding *R* values (Eqn S2) after 0.04% TX100 treatment. Each bar represents data from 60-90 cells from 2-3 independent experiments. Right panels show box plots of raw fluorescence values of ~30 cells for each of –/+ TX100 conditions from a representative experiment. C-D) FRAP results: Left panels show representative raw fluorescence recovery curves (circle) and corresponding fits (solid line, Eqn S1) for specified probe under – Ag (red) and + Ag (black) conditions. Right panels show box plots of fitted values of recovery time and mobile fraction, respectively, from multiple cells (Eqn 1 defines these parameters). Number of cells for Lyn-EGFP: 19 (– Ag) and 15 (+ Ag); and for S15-Lyn-EGFP: 19 (– Ag) and 18 (+ Ag). Box plot parameters are described in legend to Figure S1.

### Figure S8

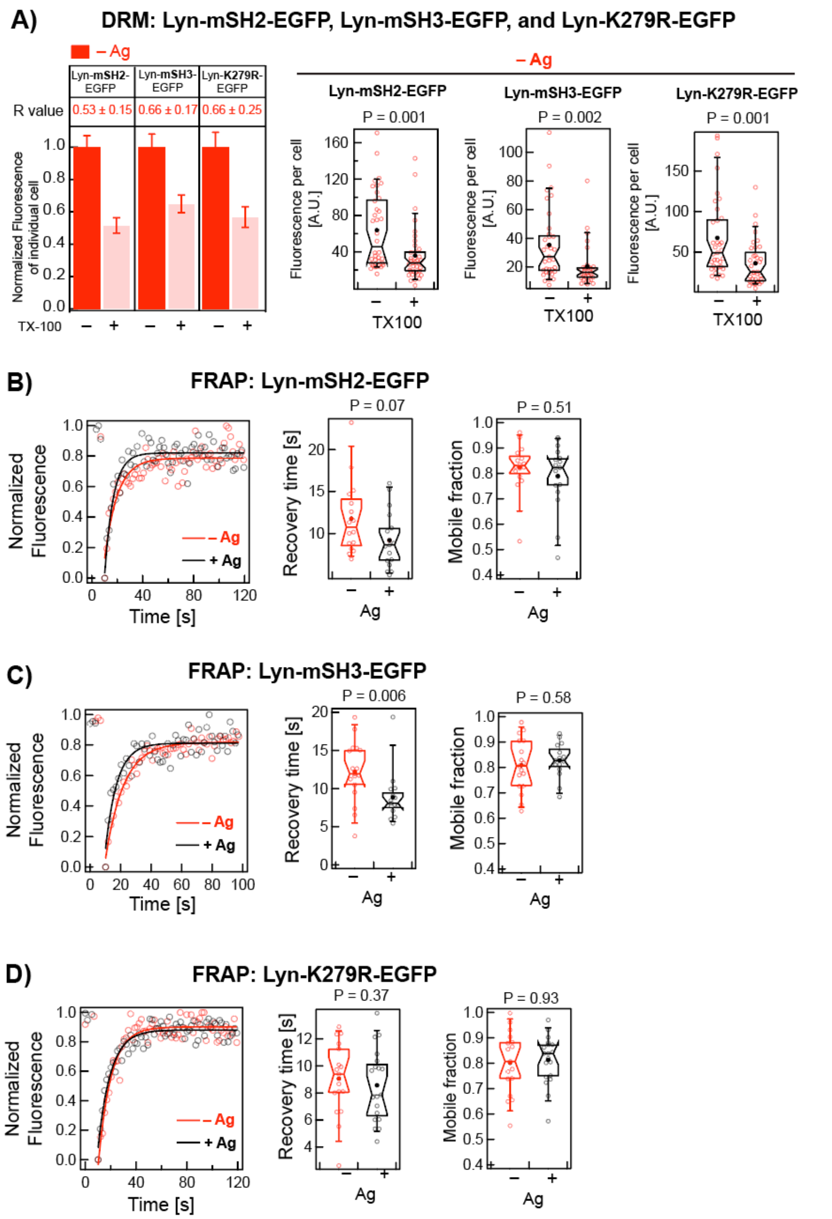


**Figure S8.** DRM and FRAP results for Lyn variants Lyn-mSH2-EGFP, Lyn-mSH3-EGFP), and Lyn-K279R-EGFP modules. A) DRM results: Left panel shows relative loss of fluorescence and corresponding *R* value (Eqn S2) after 0.04% TX100 treatment. Each bar represents data from 60-90 cells from 2-3 independent experiments. Right panels show box plots of raw fluorescence values of ~30 cells for each of –/+ TX100 conditions from a representative experiment. B-D) FRAP results: Left panels show representative raw fluorescence recovery curves (circle) and corresponding fits (solid line, Eqn S1) of specified probe under – Ag (red) and + Ag (black) conditions. Right panels show box plots of fitted values of recovery time and mobile fraction, respectively, from multiple cells (Eqn 1 defines these parameters). Number of cells for Lyn-mSH2-EGFP: 16 (– Ag) and 17 (+ Ag); for Lyn-mSH3-EGFP: 17 (– Ag) and 15 (+ Ag); and for Lyn-K279R-EGFP: 17 (– Ag) and 19 (+ Ag). Box plot parameters are described in legend to Figure S1.

#### Figure S9

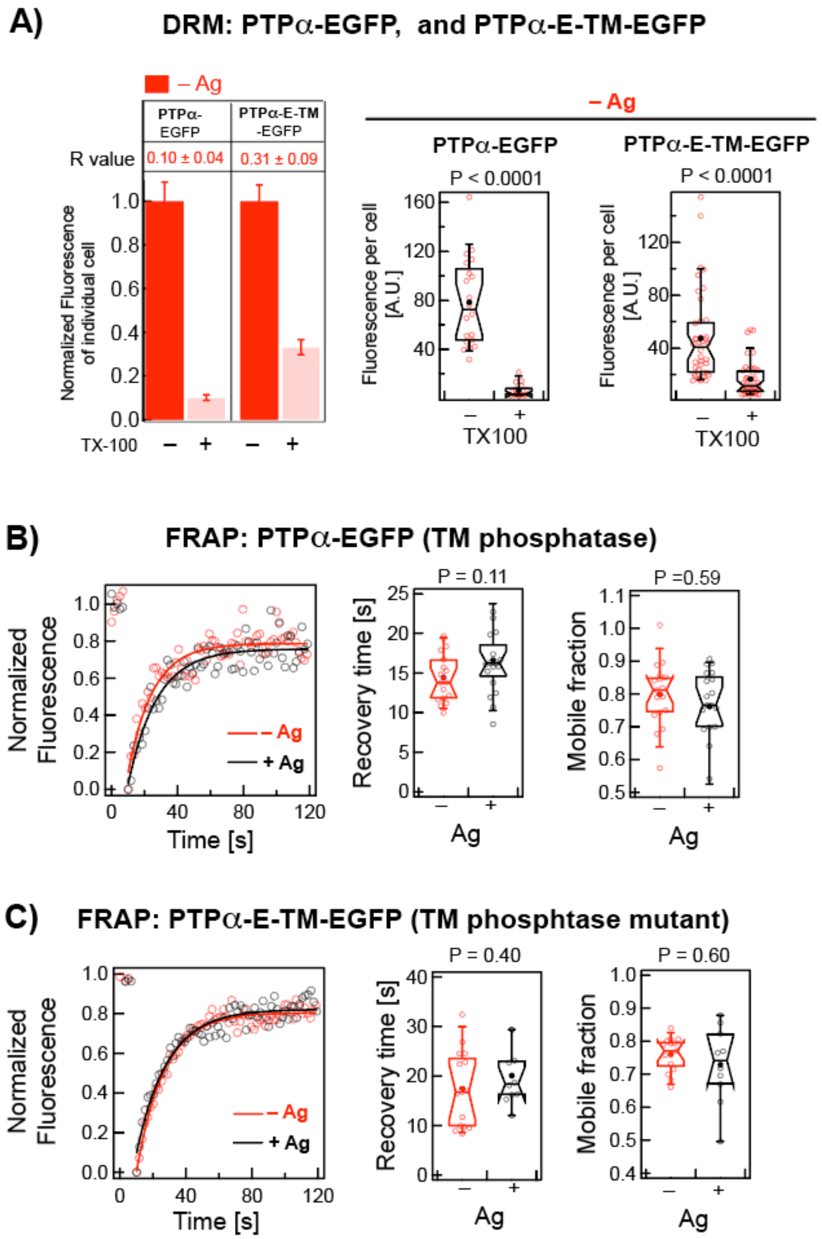


**Figure S9.** DRM and FRAP results for PTPα-EGFP and PTPα-E-TM-EGFP. A) DRM results: Left panels show the relative loss of fluorescence and corresponding *R* value (Eqn S2) upon 0.04% TX100 treatment. Each bar represents data from 60-90 cells from 2-3 independent experiments. Right panels show raw fluorescence values of ~30 cells for each of –/+ TX100 conditions from a representative experiment. B-C) FRAP results: Left panels show representative raw fluorescence recovery curves (circle) and corresponding fits (solid line, Eqn S1) of the specified probe under – Ag (red) and + Ag (black) conditions. Right panels show box plots of fitted values of recovery time and mobile fraction, respectively, from multiple cells (Eqn S1 defines these parameters). Number of cells for PTPα-EGFP: 17 (– Ag) and 19 (+ Ag); and for PTPα-E-TM-EGFP: 15 (– Ag) and 10 (+ Ag). Box plot parameters are described in legend to Figure S1.

**Table S1.** Fitting results of experimental CDFs of *D* values for all membrane probes in resting and Ag-stimulated steady-states

| Probes | Membrane association | Stimu-lation | $D_{av}$ [μm²/s][a] | $D_{fast}$ [μm²/s][b] | $D_{slow}$ [μm²/s][b] | $F_{slow}$[b] | $N_{Px}$ (No. of cells)[d] |
|---|---|---|---|---|---|---|---|
| AF488-IgE-FcεRI | TM, receptor for IgE | No[#] | 0.168 ± 0.0004 (0.167-0.168) | 0.22 ± 0.002 | 0.14 ± 0.001 | 0.71 ± 0.02 | 24582 (42) |
| | | Yes | 0.117 ± 0.0005 (0.116-0.118) | 0.16 ± 0.004 | 0.10 ± 0.002 | 0.72 ± 0.03 | 10403 (17) |
| PM-EGFP | Inner leaflet, Lo-preferring, passive lipid probe | No[#] | 0.62 ± 0.002 (0.611-0.619) | 0.68 ± 0.01 | 0.58 ± 0.006 | 0.63 ± 0.06 | 9375 (15) |
| | | Yes | 0.57 ± 0.002 (0.567-0.573) | 0.70 ± 0.03 | 0.53 ± 0.009 | 0.79 ± 0.07 | 9375 (15) |
| EGFP-GG | Inner leaflet, Ld-preferring, passive lipid probe | No[#] | 0.64 ± 0.002 (0.634-0.643) | 0.61 ± 0.004 | 0.66 ± 0.005 | 0.43 ± 0.05 | 10527 (18) |
| | | Yes | 0.69 ± 0.002 (0.685-0.691) | 0.57 ± 0.01 | 0.74 ± 0.009 | 0.31 ± 0.07 | 9775 (17) |
| S15-EGFP | Inner leaflet, Ld-preferring, passive lipid probe | No | 0.90 ± 0.003 (0.894-0.904) | 0.90 ± 0.004[c] | 0.90 ± 0.004[c] | 0.50[c] | 7474 (17) |
| | | Yes | 0.92 ± 0.003 (0.911-0.922) | 0.91 ± 0.004[c] | 0.91 ± 0.004[c] | 0.50[c] | 8050 (19) |
| Lyn-EGFP | Inner leaflet, Lo-preferring, kinase | No[#] | 0.49 ± 0.001 (0.484-0.489) | 0.57 ± 0.03 | 0.44 ± 0.008 | 0.65 ± 0.13 | 10000 (16) |
| | | Yes | 0.44 ± 0.001 (0.438-0.443) | 0.50 ± 0.02 | 0.40 ± 0.02 | 0.61 ± 0.18 | 10625 (17) |
| S15-Lyn-EGFP | Ld-preferring Lyn chimera; myristoylated lipid anchor | No | 0.64 ± 0.002 (0.638-0.647) | 0.64 ± 0.002[c] | 0.64 ± 0.002[c] | 0.50[c] | 7073 (16) |
| | | Yes | 0.67 ± 0.002 (0.663-0.670) | 0.67 ± 0.01[c] | 0.67 ± 0.01[c] | 0.50[c] | 7725 (16) |
| Lyn-mSH2-EGFP | Lyn point mutant; Arg to Ala at position 135 of SH2 module | No | 0.48 ± 0.002 (0.480-0.486) | 0.48 ± 0.002[c] | 0.48 ± 0.002[c] | 0.50[c] | 9946 (16) |
| | | Yes | 0.51 ± 0.001 (0.509-0.515) | 0.57 ± 0.01 | 0.45 ± 0.008 | 0.48 ± 0.07 | 8125 (13) |
| Lyn-mSH3-EGFP | Lyn point mutant; Trp to Ala at position 78 of SH3 module | No | 0.51 ± 0.002 (0.502-0.508) | 0.67 ± 0.04 | 0.47 ± 0.01 | 0.82 ± 0.07 | 11250 (18) |
| | | Yes | 0.49 ± 0.001 (0.487-0.492) | 0.54 ± 0.01 | 0.40 ± 0.01 | 0.38 ± 0.08 | 11778 (19) |
| Lyn-K279R-EGFP | Kinase-inactive Lyn mutant; Lys to Arg at position 279 of kinase module | No | 0.62 ± 0.002 (0.616-0.623) | 0.62 ± 0.002[c] | 0.62 ± 0.002[c] | 0.50[c] | 8827 (15) |
| | | Yes | 0.59 ± 0.002 (0.586-0.592) | 0.70 ± 0.03 | 0.55 ± 0.01 | 0.75 ± 0.07 | 9359 (14) |
| PTPα-EGFP | TM; tyrosine phosphatase | No | 0.227 ± 0.0006 (0.226-0.228) | 0.27 ± 0.007 | 0.21 ± 0.003 | 0.70 ± 0.06 | 10975 (18) |
| | | Yes | 0.199 ± 0.0005 (0.198-0.200) | 0.24 ± 0.003 | 0.18 ± 0.002 | 0.64 ± 0.03 | 12132 (20) |
| PTPα-E-TM-EGFP | TM; PTPα mutant, no cytoplasmic module | No | 0.266 ± 0.0008 (0.265-0.268) | 0.33 ± 0.007 | 0.24 ± 0.003 | 0.71 ± 0.04 | 8618 (15) |
| | | Yes | 0.240 ± 0.0006 (0.239-0.242) | 0.29 ± 0.007 | 0.22 ± 0.004 | 0.67 ± 0.06 | 8300 (14) |
| YFP-GL-GT46 | TM; passive probe | No | 0.235 ± 0.0008 (0.233-0.236) | 0.31 ± 0.007 | 0.21 ± 0.002 | 0.78 ± 0.02 | 7740 (13) |
| | | Yes | 0.211 ± 0.0005 (0.210-0.212) | 0.24 ± 0.007 | 0.19 ± 0.003 | 0.60 ± 0.02 | 9929 (16) |

[a] ± values are standard error of the mean (SEM) of the arithmetic average ($D_{av}$) of all $D$ values. Corresponding 95% confidence interval (CI) is given in parenthesis.

[b] ± values are standard deviations of the mean values obtained from the one- (Eqn A1) or two-component fitting (Eqn A2) of 30 individual bootstrapped CDFs (composed of 50% of all data each time)

[c] Values correspond to single component fit (Eqn A1); $D_{fast} = D_{slow}$, $F_{fast} = F_{slow} = 0.50$

[d] $N_{Px}$ = number of Px units from which $D$ values are determined

[#]Raw data previously published (13)

Lo = liquid ordered; Ld = liquid disordered; TM = transmembrane

PM = palmitoyl/myristoyl; GG = geranyl/geranyl; EGFP = enhanced green fluorescent protein; YFP = yellow fluorescent protein

#### APPENDIX

##### Pooling of *D* values obtained from ImFCS measurements on multiple cells and bootstrapping of raw *D* CDF followed by component analysis

The $D$ values obtained in Px units (Eqn S3) from multiple cells for a given probe at a given condition (measured on different days) are grouped to create their respective distribution. First, cumulative frequencies for each $D$ value were determined in ascending order, which were then plotted against corresponding $D$ values to generate normalized cumulative distribution function (CDF) of $D$ values using Igor Pro (Version 8; WaveMetrics, OR, USA). This CDF was fitted with the following models (Eqns A1 and A2 for one- and two-component Normal distribution models, respectively) (13).

$$\mathrm{CDF}(D) = \frac{1}{2}\left(1 + \mathrm{erf}\left(\frac{D - \mu_1}{\sigma_1\sqrt{2}}\right)\right) \tag{A1}$$

$$\mathrm{CDF}(D) = \frac{1}{2}\left[\mathrm{F}_1\left(1 + \mathrm{erf}\left(\frac{D - \mu_1}{\sigma_1\sqrt{2}}\right)\right) + (1 - \mathrm{F}_1)\left(1 + \mathrm{erf}\left(\frac{D - \mu_2}{\sigma_2\sqrt{2}}\right)\right)\right] \tag{A2}$$

In the above equations, $\mu_1$ and $\sigma_1$ are the mean and standard deviation of the first component while $\mu_2$ and $\sigma_2$ are the mean and standard deviation of the second component, $F_1$ is the fraction of first component and (1- $F_1$) is the fraction of the second component of the $D$ CDF. The best fitting model (Eqn A1 or A2) were chosen based on the absence of periodicity in the fitting residual plot and minimum reduced chi-squared value (13). A three-component model did not improve the quality of fitting in any case and therefore was not considered.

For two-component CDF fit, the component with smaller mean value, i.e., min [$\mu_1$, $\mu_2$] = $D_{slow}$ while the other component, i.e., max [$\mu_1$, $\mu_2$] = $D_{fast}$. In this case, $F_{slow}$ corresponds to the fraction of total Px units corresponding to the $D_{slow}$ component and (1-$F_{slow}$) is the fraction of Px units corresponding to the $D_{fast}$ component. For one-component CDF fit, $D_{slow}$ = $D_{fast}$ and $F_{slow}$ = $F_{fast}$ = 0.5.

*Bootstrapping of raw D values rules out eliminates possible skewing of the D distribution due to outlier Px units*

We previously showed that experimental *D* CDFs for membrane probes are often satisfactorily fitted with two Gaussian components with close values of $D_{slow}$ and $D_{fast}$ and overlapping standard deviations (13). Therein we further demonstrated that our data statistics (~10,000 *D* values) is sufficient to distinguish 10% difference between $D_{fast}$ and $D_{slow}$, i.e., $D_{fast}/D_{slow} \geq 1.1$ can be distinguished by ImFCS. Since the large set of data came from multiple cells measured on different days, we here decided to check whether the CDF of a randomly selected subset of the data represents the CDF or the entire data and whether the parameters obtained from fitting the CDFs (i.e., $D_{fast}$, $D_{slow}$, and $F_{slow}$) are statistically reliable. For this test, we chose the pooled *D* values (10,527 *D* values) of EGFP-GG in untreated cells for which we obtained the smallest difference between $D_{fast}$ and $D_{slow}$ (13). We first bootstrapped 30 times with 5% of all data each time (i.e, Number of data points per bootstrapping ($N_{BS}$) = 500). The individual bootstrapped data distributions do not show statistically significant differences among them (P > 0.1; Mann-Whitney test). The arithmetic average values of individual bootstrapped data sets ($D_{av}$) are very close (95% confidence interval range: 0.638 – 0.642 μm$^2$/s) (Figure A1). For comparison, the arithmetic average of pooled data ($D_{av,pooled}$) is 0.64 ± 0.002 μm$^2$/s (number of data points = 10,527) (13) (Table S1).

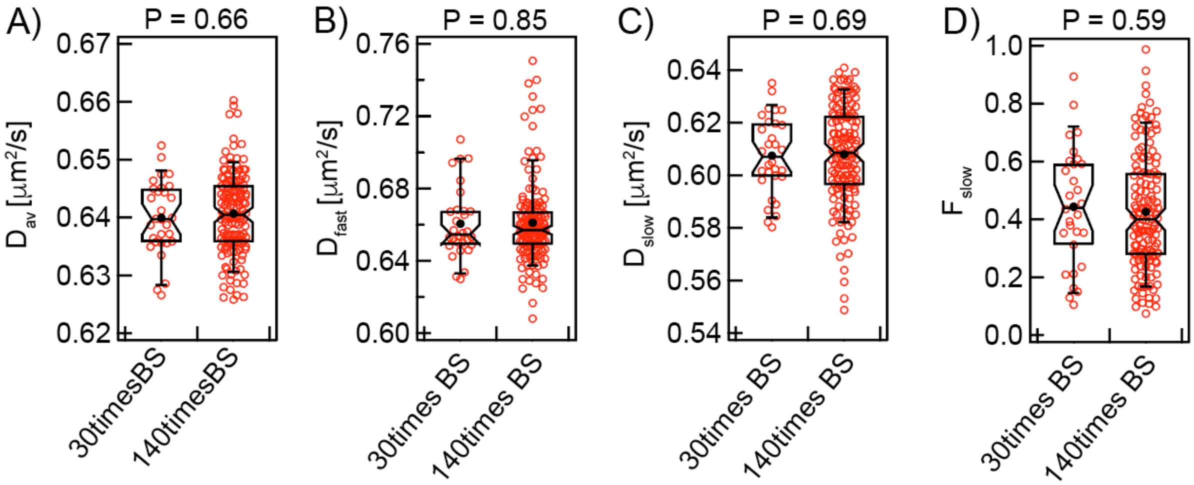


**Figure A1.** The range of values obtained from fitting bootstrapped CDFs does not become narrower with the number of bootstrapping trials (30 vs 140 times). 5% of all *D* values ($N_{BS}$ = 500) of EGFP-GG was used for each bootstrapping trial. The range of $D_{av}$, $D_{fast}$, $D_{slow}$, and $F_{slow}$ values are shown as box plots. Box height corresponds to 25th to 75th percentile and error bars represent 9th to 91st percentile of entire data set. Mean and median values are represented as solid circle and bar, respectively, located inside the box. The notches signify 95% confidence interval of the median. The statistical analysis was performed using Mann-Whitney test.

*Bootstrapped CDFs represent same underlying distribution (one-component or two-component Gaussian distributions) as that of CDF from entire data set*

The CDFs of the bootstrapped data sets ($N_{BS}$ = 500, red, Figure A2A) show little deviation from each other, and they are distributed around the CDF of all pooled data (black, Figure A2A). As expected, the spread of bootstrapped CDFs reduces as more data points are used to create these (Figure A2A-C). These narrowly distributed CDFs allowed us to do further statistical analyses to determine the underlying components with high precision.

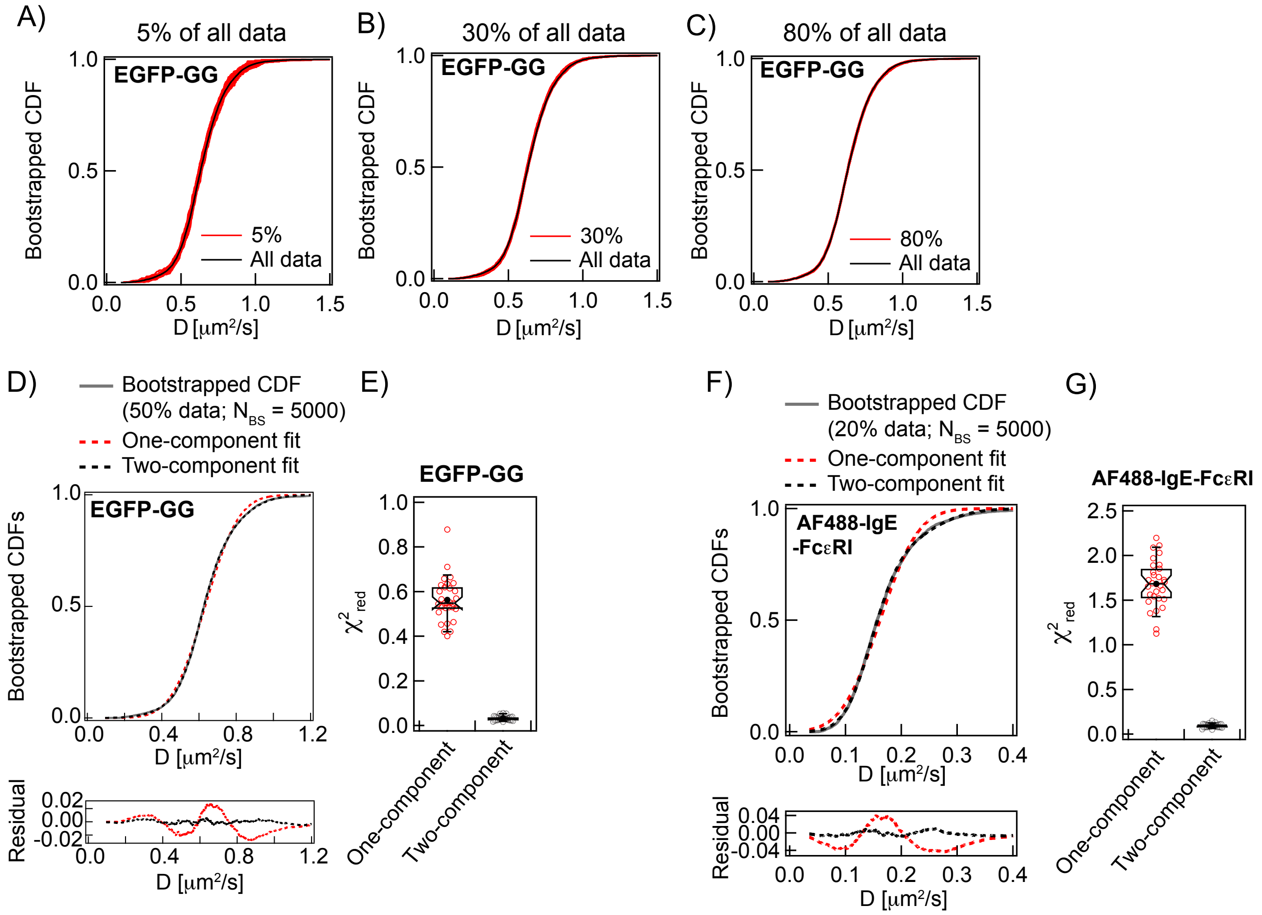


**Figure A2.** Two-component CDF model satisfactorily fits the bootstrapped data for EGFP-GG and AF488-IgE-FcεRI. **A-C**) 30 bootstrapped CDFs for EGFP-GG (red) with different $N_{BS}$ = 500 (5%), or = 3,000 (30%), or = 8,000 (80%) are separately overlaid on the single CDF (black) constructed from all 10,527 $D$ values. **D)** Fitting and residual of a representative bootstrapped CDF of EGFP-GG ($N_{BS}$ = 5000 corresponding to 50% of all data; grey solid line) with one-component model (red dotted line) and two-component model (black dotted line). **E)** Box plots of reduced chi squared ($\chi^2_{red}$) values obtained from one-component and two-component fits of all 30 bootstrapped CDFs for EGFP-GG ($N_{BS}$ = 5000 each time). **F)** Fitting and residual of a representative bootstrapped CDF of AF488-IgE-FcεRI ($N_{BS}$ = 5000 corresponding to 20% of all data; grey solid line) with one-component model (red dotted line) and two-component model (black dotted line). **E)** Box plots of reduced chi squared ($\chi^2_{red}$) values obtained from one-component and two-component fits of all 30 bootstrapped CDFs for AF488-IgE-FcεRI ($N_{BS}$ = 5000 each time). Box plot parameters are described in legend to Figure A1.

We previously showed that the CDF obtained from entire set of EGFP-GG $D$ values is satisfactorily fitted with two-component Gaussian model (13). To evaluate bootstrapped CDFs we fitted with one-component or two-component Gaussian models (Eqns A1 and A2). Figure A2D shows a representative CDF obtained from bootstrapping of EGFR-GG $D$ values with $N_{BS}$ = 5000 (corresponding to 50% of all data). The residuals plot clearly indicates that two-component model is the better model. Repetition of these analyses on all 30 bootstrapped CDFs give same conclusion which is also evident from the ~10 time lower $\chi^2_{red}$ values obtained for two-component fit compared to one-component fit (Figure A2E). We also tested the same set of analysis on the data for AF488-IgE-FcεRI for which we measured ~25000 *D* values. We previously showed the CDF of all *D* values is fit with a two-component model (13). As shown in Figure A2F-G, the bootstrapped CDFs also fit better with two-component model than one-component model. Notably, we used in this case only 20% of all data for bootstrapping. In the following section, we demonstrate the optimal data statistics required for CDF fitting with Eqns A1 and A2.

*CDFs of bootstrapped D values with 50% of all data yield high precision of the fitted $D_{fast}$, $D_{slow}$, and $F_{slow}$ values*

We fit 30 bootstrapped CDFs of EGFP-GG ($N_{BS}$ = 500 corresponding to 5% data) individually with two-component model (Eqn A2) for component analyses and resulting range of values for $D_{fast}$, $D_{slow}$ and $F_{slow}$ are given in Figure A3A-C. For reference, the $D_{fast}$, $D_{slow}$ and $F_{slow}$ values obtained after fitting the CDF of the all pooled data (*N* = 10,527) are 0.66, 0.61 and 0.41 respectively (13) (Table S1). As shown in Table A1, we obtain very close average values of all three parameters from the fitting of bootstrapped CDFs. However, while the distribution of $D_{fast}$ and $D_{slow}$ obtained from the fitting is narrow that of $F_{slow}$ is more widely distributed. The range of the values (minimum and maximum), average and standard deviation of these parameters are given in Table A1**.** We observed that increasing the number of bootstrapping trials to 140 does not narrow the range of values (Figure A4 and Table A1). Therefore, we decided to bootstrap 30 times for all our following analyses.

**Table A1.** Fitting results of bootstrapped CDFs with two-component model (Eqn A2)

| Number of bootstrapping ($N_{BS}$ = 500) | Fitted parameters of bootstrapped CDFs | Minimum value | Maximum value | Average | Standard deviation | 95% confidence interval range |
|---|---|---|---|---|---|---|
| 30 | $D_{fast}$ [μm$^2$/s] | 0.63 | 0.71 | 0.66 | 0.02 | 0.653-0.668 |
| 140 | | 0.61 | 0.75 | 0.66 | 0.02 | 0.658-0.665 |
| 30 | $D_{slow}$ [μm$^2$/s] | 0.58 | 0.64 | 0.61 | 0.01 | 0.602-0.613 |
| 140 | | 0.55 | 0.64 | 0.61 | 0.02 | 0.605-0.611 |
| 30 | $F_{slow}$ | 0.11 | 0.89 | 0.44 | 0.20 | 0.36-0.52 |
| 140 | | 0.07 | 0.99 | 0.43 | 0.20 | 0.39-0.46 |

We next resampled the data by bootstrapping (30 times) with increasing $N_{BS}$, number of *D* values per bootstrapping trial. We bootstrapped with 5% ($N_{BS}$ = 500), 15% ($N_{BS}$ = 1,500), 30% ($N_{BS}$ = 3,000), 50% ($N_{BS}$ = 5,000) and 80% ($N_{BS}$ = 8,000) of all EGFP-GG *D* values (10,527, Table S1), and we constructed CDFs for each case. Box plots of fit parameters for these bootstrapped CDFs using the two-component model are given in Figure A3A-C. The range of $D_{fast}$, $D_{slow}$ and $F_{slow}$ values becomes narrower with larger $N_{BS}$. However, as shown in Figure A3D-F, the Mann-Whitney test comparing a given parameter (e.g., $D_{slow}$) between any two bootstrapped CDFs constructed from different $N_{BS}$ (e.g., 5% and 80% of all data) show P value > 0.05. This comparison indicates that for most parameters the CDFs are not statistically different across this broad range of $N_{BS}$. The only exception is $F_{slow}$ obtained from the CDFs with 30% and 50% of all data.

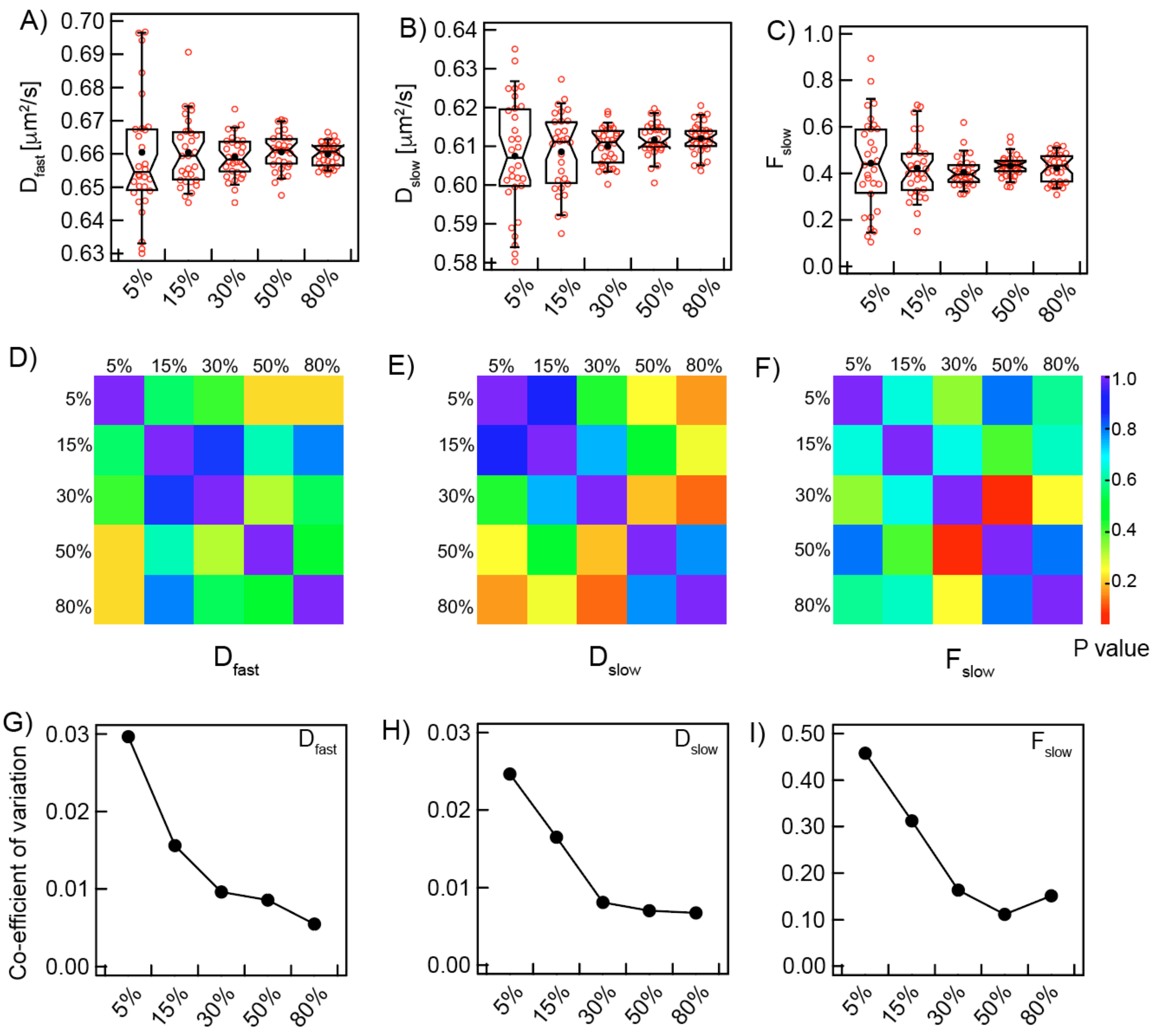


**Figure A3.** The precision of $D_{fast}$, $D_{slow}$, and $F_{slow}$ increases with the number of data points ($N_{BS}$) used to create the bootstrapped CDFs. *D* values of EGFP-GG in resting cells (- Ag) (Table S1) used in this example. 30 bootstrapped CDFs with different $N_{BS}$ (500 (5%), 1,500 (15%), 3,000 (30%), 5,000 (50%), and 8,000 (80%) of a total of 10,527 *D* values) were individually fitted with two-component model (Eqn A2). **A-C)** 30 fitted values for each parameter from CDFs obtained after bootstrapping with increasing $N_{BS}$. **D-E)** P values (nonparametric Mann-Whitney test) for each pair of sets of fitted data for $D_{fast}$, $D_{slow}$, and $F_{slow}$ obtained from CDFs with different $N_{BS}$. **G-I)** Precision of $D_{slow}$, $D_{fast}$ and $F_{slow}$, as quantified from their respective coefficient of variation, as a function of $N_{BS}$.

Figure A3A-C shows that as the range of $D_{fast}$, $D_{slow}$, and $F_{slow}$ values becomes narrower with larger $N_{BS}$, estimation of these parameters becomes more precise. High precision is necessary to compare subtle changes of a given parameter between conditions (e.g., the change of $D_{slow}$ of EGFP-GG before and after stimulation with Ag). We found that 5% of all data was sufficient to achieve highly precise estimations of $D_{fast}$ and $D_{slow}$ as determined from their respective coefficient of variation (CoV) plots (Figure A3G-I): CoV < 0.1 for both parameters. For $F_{slow}$, we needed at least 30% of all data ($N_{BS}$ = 3,000) to achieve CoV ~ 0.1. To be conservative, we use 50% data points in ImFCS studies presented in the main text to achieve highly precise estimations of $D_{fast}$, $D_{slow}$ and $F_{slow}$ based on our data sets and analyses.

As a final check for bias, we performed CDF analyses on five different sets of 30 bootstrapping trials using $N_{BS}$ = 5000 and the EGFP-GG data set. As illustrated in Figure A4, the five different bootstrapping sets yielded same distributions for the each of $D_{fast}$, $D_{slow}$, and $F_{slow}$.

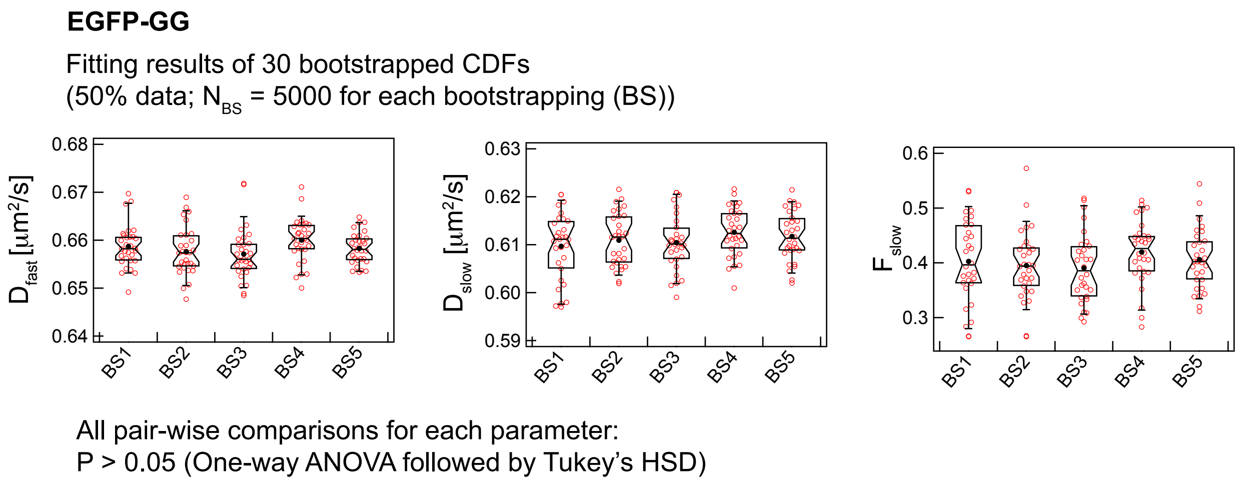


**Figure A4.** Two-component fitting of multiple sets of 30 bootstrapped CDFs yield statistically indistinguishable distribution for each of $D_{fast}$, $D_{slow}$, and $F_{slow}$ parameters.
